# A broadly-neutralizing antibody against *Ebolavirus* glycoprotein that potentiates the breadth and neutralization potency of other antibodies

**DOI:** 10.1101/2024.06.21.600001

**Authors:** Francesca R. Donnellan, Vamseedhar Rayaprolu, Pramila Rijal, Victoria O’Dowd, Amar Parvate, Heather Callaway, Chitra Hariharan, Dipti Parekh, Sean Hui, Kelly Shaffer, Ruben Diaz Avalos, Kathryn Hastie, Lisa Schimanski, Helena Müller-Kräuter, Thomas Strecker, Ariane Balaram, Peter Halfmann, Erica Ollmann Saphire, Daniel J. Lightwood, Alain R. Townsend, Simon J. Draper

## Abstract

Ebolavirus disease (EVD) is caused by multiple species of *Ebolavirus*. Monoclonal antibodies (mAbs) against the virus glycoprotein (GP) are the only class of therapeutic approved for treatment of EVD caused by *Zaire ebolavirus* (EBOV). Therefore, mAbs targeting multiple *Ebolavirus* species may represent the next generation of EVD therapeutics. Broadly reactive anti-GP mAbs were produced; among these, mAbs 11886 and 11883 were broadly neutralizing *in vitro*. A 3.0 Å cryo-electron microscopy structure of EBOV GP bound to both mAbs shows that 11886 binds a novel epitope bridging the glycan cap (GC), 3_10_ pocket and GP2 N-terminus, whereas 11883 binds the receptor binding region (RBR) and GC. *In vitro*, 11886 synergized with a range of mAbs with epitope specificities spanning the RBR/GC, including 11883. Notably, 11886 increased the breadth of neutralization by partner mAbs against different *Ebolavirus* species. These data provide a strategic route to design improved mAb-based next-generation EVD therapeutics.

## Introduction

*Zaire ebolavirus* (EBOV) and *Sudan ebolavirus* (SUDV) were both first described in 1976; since then multiple species of *Ebolavirus* have spilled over from animal reservoirs to cause fatal *Ebolavirus* disease (EVD) in humans ^1^. Subsequently, the emergence of *Tai Forest ebolavirus* (TAFV) ^2^ and *Bundibugyo ebolavirus* (BDBV) ^3^, which also cause EVD in humans, and the discovery of *Reston* (RESTV) and *Bombali ebolaviruses* (BOMV) in pigs and bats respectively ^4,5^, has further illustrated the risk of outbreaks of EVD caused by known and yet to emerge *Ebolaviruses* from reservoir species. The large-scale outbreak in West Africa in 2013-2016, which caused >28,000 cases and claimed over 11,000 lives, brought Ebola to global attention, and accelerated the development of vaccines and therapeutics. The result was that a vaccine for use in outbreaks was prequalified by the World Health Organization in 2019 ^6^, and two monoclonal antibody (mAb)-based therapeutics to treat EVD were approved by the US Food and Drug Administration in 2020 ^7,8^. However, despite this progress, there have been nearly annual outbreaks of EVD since 2017, with over 2,000 lives claimed ^9^. Moreover, ensuring vaccine coverage, risk of vaccine breakthrough and recrudescence of virus in survivors all present on-going challenges to preventing outbreaks alongside those posed by ever increasing human interactions with animal reservoirs ^10^.

The only therapeutics approved for use against EVD are both mAb formulations: Ebanga^TM^ (also called Ansuvimab-zykl, a single antibody called “mAb114”) ^7^ and Inmazeb^TM^ (also called REGN-EB3, a cocktail of three antibodies named “REGN3479, REGN3471 and REGN3470”)^8^. Whilst these therapeutics represent a step change in our ability to treat EVD and undoubtedly have already saved lives, each of these had limited efficacy in field trials: approximately one third of patients in Ebanga or Inmazeb treated groups died when these drugs were trialled in an outbreak of EVD caused by EBOV ^11^. Efficacy was reduced for patients with higher viral loads or a longer interval between onset of symptoms and treatment ^11^. In addition, there is an expectation of further reduced efficacy against EVD caused by other *Ebolavirus* species ^12^. Therefore, there is an ongoing effort to develop a robust pipeline of additional candidate therapeutics or treatment strategies with increased potency and breadth of action against multiple species of *Ebolavirus* ^10^. This is required if we are to meet the ongoing challenges presented by *Ebolaviruses* and to contribute to preparedness for future outbreaks.

Therapeutic mAbs, including those that constitute Inmazeb and Ebanga, target the EBOV glycoprotein (GP). The GP is the only transmembrane *Ebolavirus* surface protein and is responsible for entry of virus into host cells; mediating cell attachment, host receptor binding, viral and host cell membrane fusion, as well as having roles in viral budding and release. Precursor GP is cleaved by host furin proteases to yield GP1 and GP2 disulphide-bonded heterodimers which non-covalently associate into a chalice-shaped trimer ^13,14^. GP1 (residues 30-501) contains a core domain, the receptor binding region (RBR), glycan cap domain (GC) and mucin-like domain (MLD). In the pre-fusion conformation, GP1 forms the bowl of the chalice, with the RBR largely occluded by the GC and MLD, the latter being heavily glycosylated and variable between species ^15^. GP2 (residues 502-676) contains the internal fusion loop (IFL), heptad repeats (HR1 and HR2), membrane proximal external region (MPER), the transmembrane domain (TM) and cytoplasmic tail ^14^. GP2 cradles the GP1 chalice and forms the stalk that anchors the GP to the viral membrane, with the IFL extending around the neighbouring GP1-GP2 heterodimer. During the infection cycle, the virus is internalized into the cellular endosomal pathway via macropinocytosis, whereupon the GC and MLD domains are sequentially cleaved by host cathepsins to form GP_CL_ with an exposed RBR ^16,17^. Upon binding to the host receptor, Niemann-Pick C1 (NPC1), and additional triggers, the GP undergoes substantial rearrangements to form a classic six-helix bundle that mediates membrane fusion ^18^.

In recent years, a number of broadly reactive candidate mAbs have been developed and tested in animal challenge models against EBOV, SUDV and BDBV; several with encouraging broad efficacy data. Whilst these mAbs have been more thoroughly reviewed elsewhere ^19,20^, several key themes are emerging. Firstly, despite the diversity of the MLD and overall low sequence conservation between GP from different *Ebolavirus* species, there are key conserved sites of vulnerability on the GP, including the IFL, the 3_10_ pocket of the GP1 core and the RBR, that can be targeted by broadly reactive mAbs. Secondly, despite a high degree of conservation in these specific sites across species, antibodies targeting these epitopes are still able to select for infection competent viral escape mutants *in vitro* and in animal challenge models ^21–25^. However, more encouragingly, a recent study of the Inmazeb cocktail demonstrated the power of combining three mAbs targeting non-overlapping sites in the GP into a cocktail to mitigate selection for EBOV escape ^26^. There is, therefore, broad consensus that it is relevant to consider how individual candidate mAbs will fit into cocktail combinations for therapeutic development. Finally, the power of combining mAbs into cocktails may not be limited to only reducing risk of viral escape, but also to benefit from co-operative effects between partner mAbs to enhance potency. This potential for mAb candidates to synergize may thus help address the need for increased potency that is required of the next generation of mAb therapeutics against EVD ^12,21,27,28^.

Here, we report a novel mAb against the GP with a tripartite epitope extending down the outside of the GP chalice, that is not only broadly neutralizing, but also broadly potentiating – enhancing the potency and neutralization breadth of multiple antibodies (with different epitopes spanning the RBR and GC) when tested in combination.

## Results

### Isolation of broadly reactive monoclonal antibodies against *Ebolavirus* GP

To generate anti-*Ebolavirus* GP mAbs, a New Zealand white rabbit was immunized with a mixture of RAB-9 cells expressing EBOV GP, SUDV GP, BDBV GP and TAFV GP. B cells were cultured and antigen-specific IgG secreting cells identified using UCB Pharma’s single B cell antibody discovery platform. IgG variable region sequences were recovered from single cells and expressed as recombinant mAbs. Binding of recombinant mAbs to full-length transmembrane GP expressed on the surface of cells was confirmed and a panel of five rabbit IgGs identified (11883, 11886, 11889 11892, and 11897) that were cross-reactive to GPs from all three species of *Ebolavirus* that have caused outbreaks of EVD in humans (**Figure 1A**). IgGs in the panel were confirmed as distinct clones by sequencing. In addition, binding to GP from TAFV, BOMV and RESTV, the remaining known *Ebolavirus* species, was evaluated. 11897 did not bind BOMV or RESTV, but the other four mAbs recognized all three additional viral GPs (**Figure S1A**).

**Figure 1.**
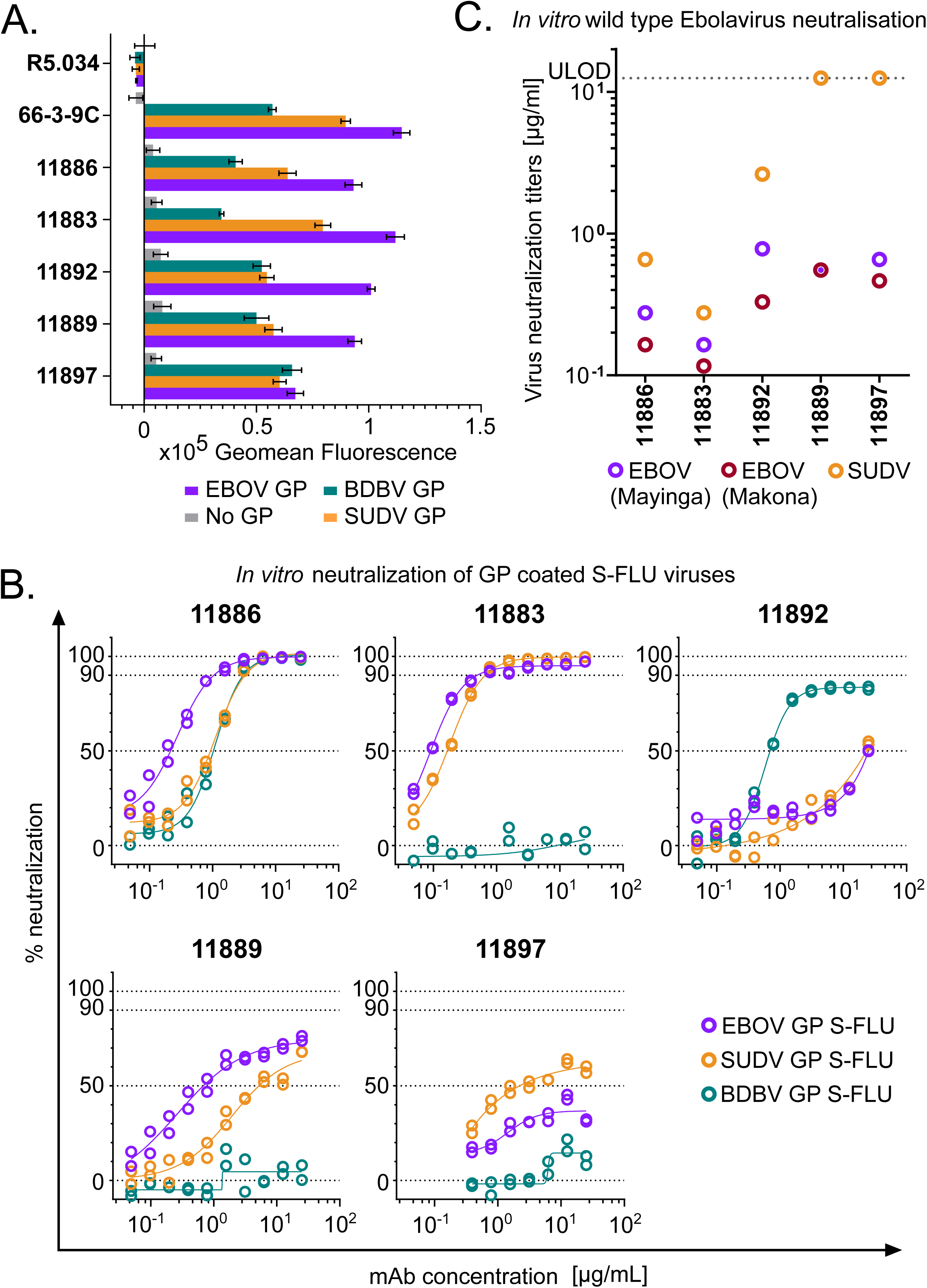
Binding to GP and virus neutralization by rabbit mAb panel. **A) Binding of broadly reactive rabbit mAb panel to full-length transmembrane *Ebolavirus* GPs.** MDCK SIAT-1 cells expressing EBOV GP (purple), SUDV GP (orange), BDBV GP (teal) or parental cells not expressing GP (No GP, grey) were incubated with 20 μg/mL of mAb, and mAb binding detected using an Alexa Fluor 647 anti-IgG conjugate secondary antibody. R5.034 (non-anti-GP mAb) and 66-3-9C (broadly reactive anti-GP mAb) were included as negative and positive controls, respectively. Background (mean of 8 wells per assay plate incubated with relevant secondary only) was subtracted from data. Mean and SEM of duplicates shown for each mAb. **B) *In vitro* neutralization of *Ebolavirus* GP pseudotyped S-FLU viruses.** S-FLU viruses coated with EBOV GP (purple) (NP_066246.1), SUDV GP (orange) (YP_138523.1) and BDBV GP (teal) (YP_003815435.1) were incubated with mAb, and then MDCK SIAT-1 cells were infected with the mAb-virus mixture. Percent neutralization of S-FLU virus by titrated mAb was calculated using maximal (no antibody added) and minimal fluorescence (no virus added) signals within each assay. Duplicates within assay for each mAb test concentration and calculated non-linear regression curves are shown. **C) *In vitro* neutralization of wild-type *Ebolaviruses*.** EBOV Mayinga (purple) (GenBank accession: AF086833), EBOV Makona (maroon) (GenBank accession: KJ660347) or SUDV (orange) (GenBank accession: FJ968794) were incubated with test mAb, and Vero E6 cells then infected with the mAb-virus mixture. Virus neutralization titers (VNT) indicate the lowest concentration of mAb at which inhibition of the cytopathic effect on infected Vero E6 cells was observed. Lower limit of detection = 0.006 μg/mL. Upper limit of detection (ULOD) = 12.5 μg/mL.

### 11886 is a broadly neutralizing anti-GP mAb *in vitro*

The panel of five novel rabbit mAbs were next tested for their ability to neutralize GP pseudotyped viruses (**Figure 1B, S1B**) and wild-type *Ebolaviruses* (**Figure 1C**) *in vitro*. S-FLU viruses with a GFP reporter can be successfully pseudotyped with GP from EBOV, SUDV and BDBV and handled at BSL-2 ^29^. S-FLU viruses could not be successfully pseudotyped with BOMV or TAFV GP. The lack of success of BOMV pseudotyping (due to a difference in the receptor binding site which limits infection of assay target cells) has previously been observed with a lentivirus system and may explain the lack of infection of MDCK SIAT cells in the S-FLU platform with this BOMV GP sequence ^30^. Neutralization of wild type BDBV was not tested. Despite all mAbs in the panel being able to bind to EBOV, SUDV and BDBV GPs, at the highest concentrations tested in the neutralization assay, only mAb 11886 was able to fully neutralize all three GP coated S-FLU viruses. 11883 fully neutralized EBOV and SUDV GP coated S-FLU viruses, whilst at the same concentrations was unable to even partially neutralize BDBV GP S-FLU. Encouragingly, both 11886 and 11883 individually also potently inhibited the cytopathic effect (CPE) of wild-type EBOV and SUDV on Vero cells, with inhibition of CPE of all three tested isolates seen at concentrations <1 μg/mL. Neutralization of RESTV GP S-FLU was also tested, with both 11886 and 11883 showing potent inhibition (**Figure S1B**). 11892 displayed partial neutralization of EBOV and SUDV GP coated S-FLU viruses at 12.5 μg/mL, as well as being able to neutralize BDBV GP S-FLU, although neutralization plateaued at ∼90 %; this mAb also reduced CPE of both wild-type EBOV and SUDV at concentrations <5 μg/mL. 11889 and 11897 showed minimal or partial neutralization of EBOV and SUDV GP S-FLU viruses, and were only able to inhibit CPE of wild-type EBOV, but not SUDV at the concentrations tested. Given the highly encouraging *in vitro* neutralization data observed with 11886 and 11883, we proceeded to map the binding sites of these novel antibody clones.

### Broadly-neutralizing anti-GP mAbs 11886 and 11883 have distinct epitopes

The novel rabbit mAbs were initially assigned to one of three broad epitope bins, (i) base, (ii) RBR/GC or (iii) IFL, by competitive binding analysis with other antibodies of known epitope (**Figure 2**). The reference antibodies c4G7, c2G4, 6541, CA45, FVM02, ADI-15946 and ADI-15878 have previously described epitopes that encompass the length of the IFL or that are present around the base of the GP chalice ^24,31–33^; whilst reference antibodies 6660, 6662, 66-3-9C and 040 have previously been assigned to either RBR or GC epitope regions in similar competition assays ^34^. Each test mAb was chemically biotinylated, then mixed with a 10-fold excess of unbiotinylated reference antibody before incubation with EBOV GP expressed on MDCK SIAT-1 cells. The amount of biotinylated antibody remaining after washing was determined using a streptavidin Alexa Fluor 647 conjugate and compared to binding in the presence of a non-GP targeting mAb control in order to determine the degree of binding competition. The five novel rabbit mAbs in this study fell into two distinct competition groups (**Figure 2**). 11883, 11889 and 11897 formed a distinct competition group that competes with anti-RBR/GC mAbs, but not those that bind the base or IFL; in contrast, 11886 and 11892 competed with each other and base-binding mAbs, but not with those that bind the GC, RBR or IFL (except for ADI-15946. As ADI-15946 only contacts the base of the IFL adjacent to the GP2 N terminus, and does not have an epitope that covers the stem or fusion peptide (FP) of the IFL (unlike ADI-15878, CA45 or FVM02), competition with only ADI-15946 and no other IFL mAbs suggests an epitope that overlaps the region adjacent to the GP2 N terminus rather than the IFL directly.

**Figure 2.**
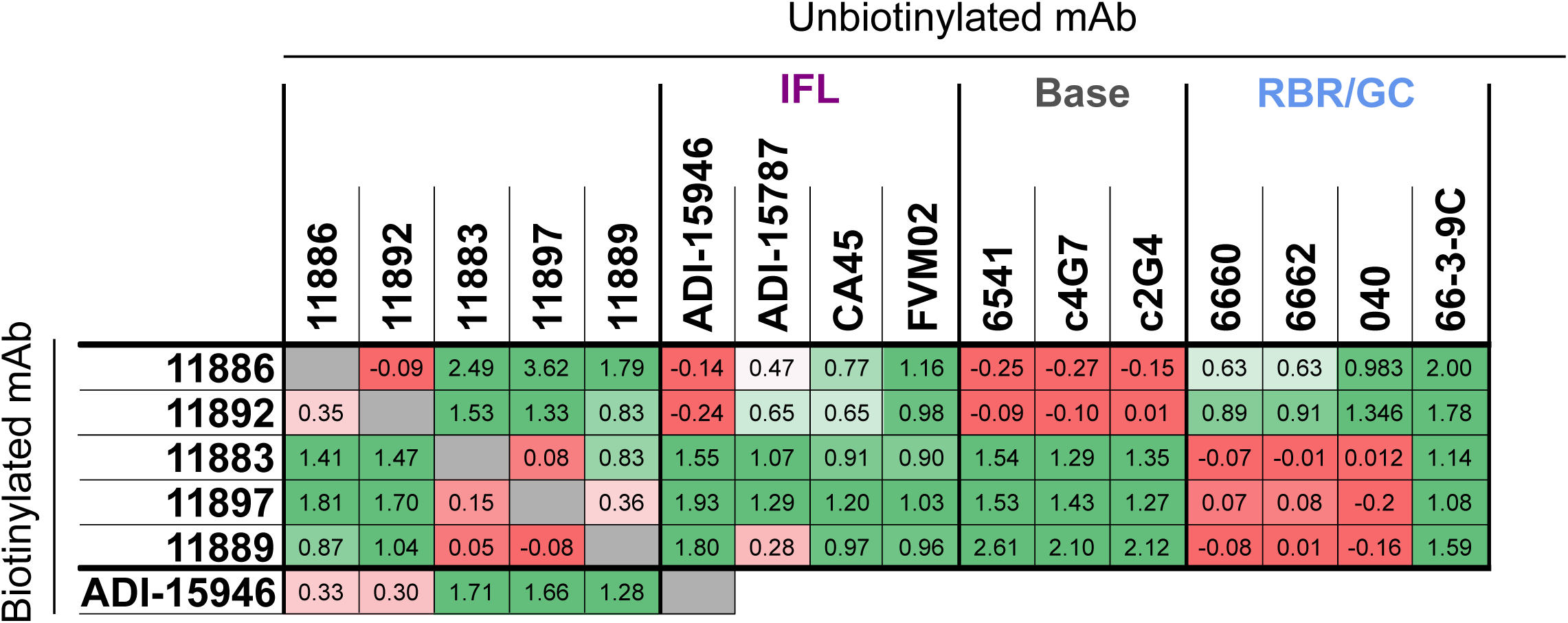
Competition of mAbs for binding to EBOV GP (Mayinga, NP_066246.1) expressed on cells. Table summarizes competition experiments between pairs of mAbs for binding to GP: 1 = signal when the biotinylated antibody is incubated with non-GP mAb or PBS (max); 0 = signal when competed against self (min). Values were calculated using the formula: (max-X)/(max-min), where X = mean signal of 6 replicate wells within each assay plate for a given competition pair. Table colored on scale from red (0; strong competition) to green (≥1; no competition). Grey; self-competition. IFL; internal fusion loop. RBR; receptor binding region. GC; glycan cap. Background (mean of 12 wells per assay plate) was subtracted from all data before calculation of competition.

Given that the IFL has previously been described as a key epitope region for broadly reactive mAbs targeting *Ebolaviruses* ^31,32,35,36^, but did not seem from our initial analysis to be the target of any of our novel rabbit mAbs we next sought to confirm this finding by an independent analysis. Notably, across *Filoviruses,* the IFL has functional and structural similarities, however whilst the IFL base, stem and FP sequences are highly conserved between *Ebolaviruses*, only the FP is relatively highly conserved in the related Filovirus, Marburg virus (MARV) (**Figure S2A**). MDCK SIAT-1 cells were therefore transduced with a DNA construct comprising the full-length EBOV GP with residues 512-555 replaced with residues 513-556 from MARV GP, i.e. this construct was designed to create a chimeric sequence comprising of the EBOV GP with the MARV GP IFL sequence (**Figure S2A**). We confirmed that i) this chimeric GP, if cleaved with thermolysin (THL) to remove the GC and MLD, bound to recombinant NPC1 (host receptor) and the pan-filovirus RBR-binding mAb MR78; and that ii) known EBOV IFL binding mAbs lost binding to the chimeric GP, whilst mAbs against other domains and mAb FVM02 (which specifically binds the conserved FP sequence ^32^) retained binding of the chimeric GP (**Figure S2B**). Notably, none of the five novel rabbit mAbs showed dependency on the presence of the EBOV IFL for binding (**Figure S2C**), despite this being one of the major conserved regions between GPs from different *Ebolavirus* species and a target of other previously described broadly reactive anti-GP mAbs.

These data were thus consistent with our previous observation and the lack of binding competition between any of the rabbit mAbs and the anti-IFL reference mAbs, except ADI-15946. Interestingly, ADI-15946 also showed greatly reduced binding to the chimeric GP, as occurred for the other anti-IFL reference mAbs (**Figure S2B**), suggesting a distinct binding mode to the rabbit mAbs 11886 and 11892. However, the epitope of ADI-15946 includes the base of the IFL as well as the N-terminus of GP2 and the 3_10_ pocket of GP1 (**Figure S4A**) ^28^, therefore the observed competition between 11886, 11892 and ADI-15946 would indicate some binding overlap is occurring in the GP base region, but not in the IFL residues swapped in the chimeric GP.

These data also indicated that the broadly-neutralizing mAbs 11886 and 11883 bind to non-competing areas of the GP, and both appeared to bind outside the IFL region as defined by competitive binding analyses with these established reference antibodies and to the chimeric GP.

### Glycan cap dependency of 11886 binding

The above binding competition data suggested that three of the mAbs, including 11883, bind in the RBR/GC on the GP molecule. To further confirm these findings, we exposed cell surface-expressed GP to THL; this enzymatic cleavage of GP sequentially deletes the MLD and GC to expose the receptor binding site, mimicking the natural action of cathepsins in target cells ^17^. THL-cleaved GP (GP_CL_), typically no longer reacts with antibodies directed against the GC or MLD (as for mAbs P6 or 040, included as a THL-digestion sensitive comparators ^34^), but does react with antibodies directed against the base of GP which remains after cleavage (as for ADI-15946 and 6541, included as comparators for this group of mAbs ^28,34^). Antibody MR78 can only bind to EBOV GP when the GC and MLD are removed exposing the RBR ^37^, and was also included as a control to confirm THL digestion of GP in the assay. Rabbit mAbs 11883, 11889 and 11897 which compete with RBR and GC binding mAbs, as expected, failed to bind EBOV GP_CL_ (**Figure 3A, S3A**) and failed to immunoprecipitate the portion of the cleaved GP corresponding to GP2 and remaining GP1 core (∼25 kDa) (**Figure 3B, S3B**). In contrast, rabbit mAb 11892 (from the base-binding competition group) showed no reduction in binding to GP_CL_ from these species (**Figure 3A, S3A**) and, like base-binding mAb 6541, could immunoprecipitate GP2 and remaining GP1 core (**Figure 3B, S3B**). 11886, however, exhibited unusual behaviour: despite previously competing for binding with base-binding mAbs suggesting an epitope away from the GC, 11886 also failed to bind EBOV GP_CL_ (**Figure 3A**). This sensitivity to the presence of the GC was shown across both the immunofluorescence and immunoprecipitation assays (**Figure 3A,B**), and was replicated for BDBV GP and TAFV GP (**Figure S3A,B**). Consistent with reduced binding, 11886 and 11883 showed a large reduction in EBOV S-FLU neutralization when the virus was pre-treated with THL (**Figure 3C**). This contrasted with comparator mAbs MR78 (receptor-binding site binder), ADI-15946 (base binder) and 2G4 (base binder) which all showed improved neutralization of the digested virus. MR78 has been reported to only neutralize EBOV when the GP is cleaved, an effect reproduced in this assay resulting in the large change in fluorescence intensity upon treatment of the virus with THL ^37^. ADI-15946 (3_10_ pocket and IFL binder in the base of the GP) which can neutralize EBOV with full-length GP has previously been shown to have enhanced neutralization of EBOV upon removal of the GC, and this effect was reproduced in this assay ^28^. The change in fluorescence intensity is smaller compared to that of MR78 as ADI-15946 can also neutralize EBOV GP S-FLU without THL treatment. 2G4, a GP-base-binding mAb from the ZMapp cocktail ^38^, was included as an additional comparator and showed a similar profile to ADI-15946 suggesting that it too modestly benefits from increased access to its epitope by removal of the GC and MLD.

**Figure 3.**
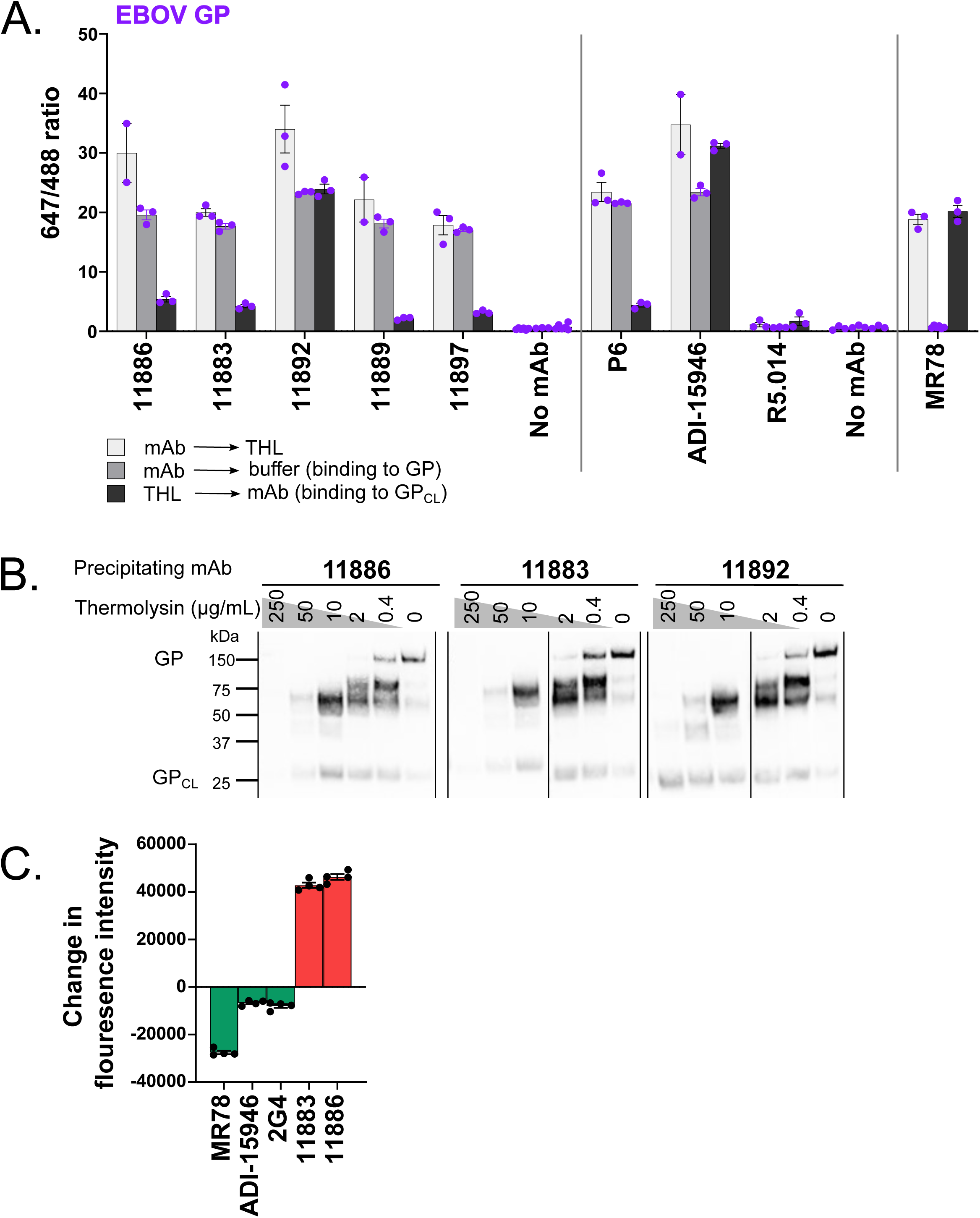
Glycan cap dependency of 11886 binding and neutralization. **(A) Antibody-mediated inhibition of thermolysin (THL) cleavage of cell surface-expressed EBOV GP was assessed in an immunofluorescence assay.** For each test mAb, binding to GP was tested under three conditions: GP-expressing cells were pre-incubated with test mAbs, then cells were treated with THL (light grey); GP-expressing cells were pre-incubated with test mAbs, then cells were treated with buffer only without enzyme (mid-grey); and GP-expressing cells were pre-incubated with THL to produce GP_CL_, then incubated with test mAbs (dark grey). mAb binding was then detected using Alexa Fluor 647 conjugate and cells stained with wheat germ agglutinin Alexa Fluor 488 conjugate. Mean and SEM for triplicate wells within the assay are shown. **(B) Immunoprecipitation of thermolysin cleaved EBOV GP by 11886 and 11883 shows loss of binding to cleaved GP compared to 11892.** GP-expressing cells were treated to biotinylate surface proteins, then incubated with increasing concentrations of THL. Cells were washed then lysed. GP was immunoprecipitated from cell lysates using Protein A Sepharose and anti-GP mAb of interest. Samples were run on reducing SDS-PAGE, and bands were revealed using streptavidin Alexa Fluor 647 conjugate. Band at ∼150 kDa is full-length GP; band at ∼25 kDa is GP_CL_ (GP1 core and GP2); other bands are products of sequential cleavage of MLD and GC. For immunoprecipitation with 11889, 11897 and 6541 see **Figure S3**. **(C) 11886 and 11883 lose ability to neutralize thermolysin-cleaved EBOV GP S-FLU pseudovirus.** The ability of mAbs to neutralize EBOV GP S-FLU virus and THL-treated EBOV GP S-FLU was tested at a single concentration of mAb. Fluorescence intensity (FI) indicates degree of infectivity of the viruses. After background correction, the change in fluorescence intensity was calculated by: FI_THL-treated virus_ – FI_Untreated virus_. mAbs that neutralize the THL-treated virus better than the untreated virus (green bars) will give negative values. mAbs that neutralize the untreated virus better than the THL-treated virus (red bars) will give positive values. Mean and SEM shown for four replicate wells within each assay.

From this epitope analysis, 11886 and 11883 therefore bind distinct epitopes that are none-the-less both reliant on the presence of the GC domain, as evidenced by a loss of binding and neutralization when the GP is cleaved. Compared to previously published broadly neutralizing mAbs that bind to the base of the GP, 11886 is unusual in that its binding to the GP is not dependent on the FL but does have GC involvement. This suggested it has an epitope that spans the GC on GP1 and GP2. We therefore sought to define the epitopes of these two most broadly neutralizing and non-competing mAbs in the panel at higher resolution using structural methods.

### 11883 binds the GC and RBR interior of the GP chalice

We next determined a high-resolution map and model of the EBOV GP ectodomain (GPdTM) in complex with both 11883 and 11886 Fabs by Cryo-EM. No symmetry was applied to the analysis in order to precisely map the epitopes of these two broadly neutralizing antibodies and investigate the unusual GC-dependency of 11886 (**Figure 4A-C**).

**Figure 4.**
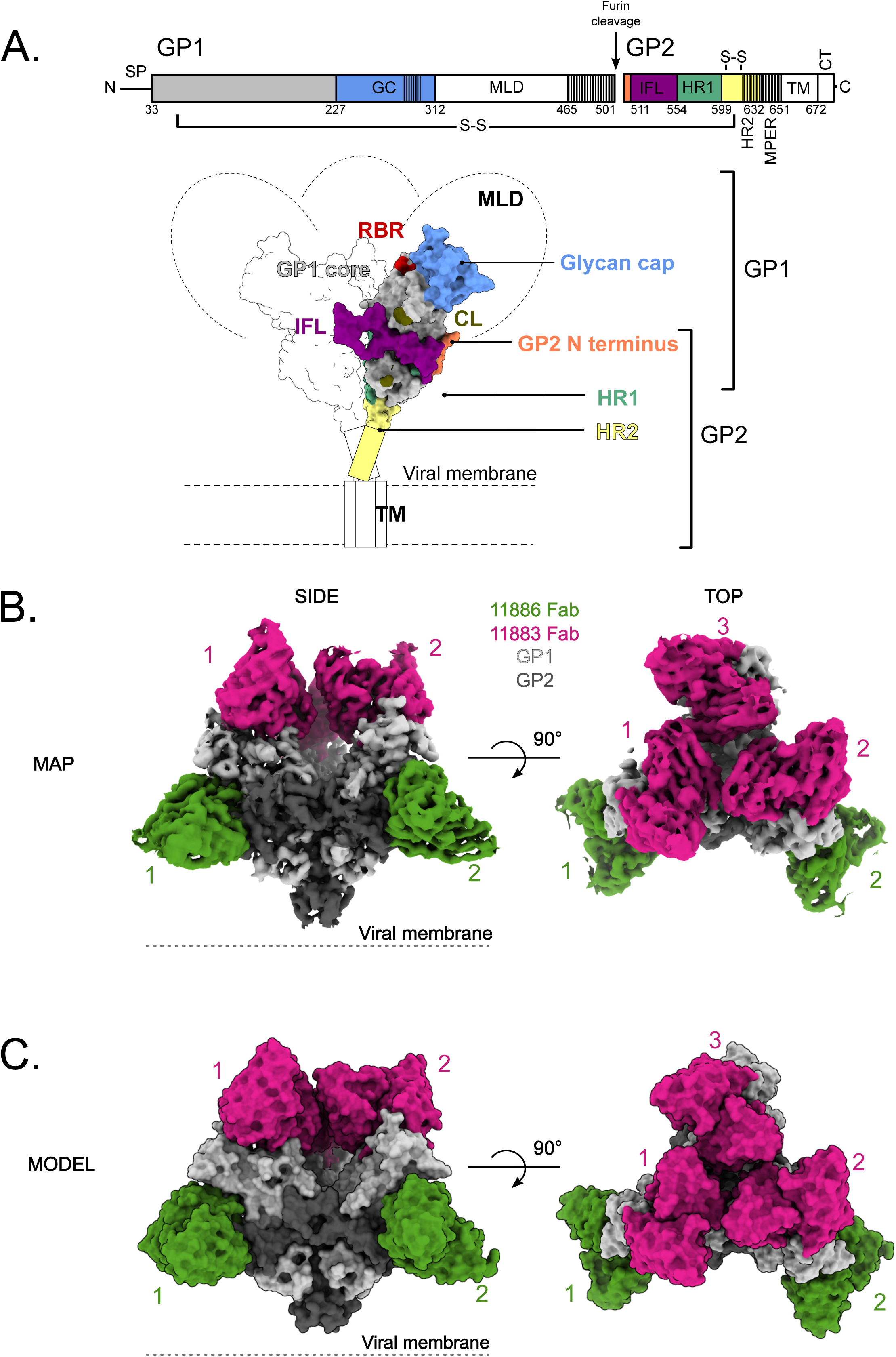
Structural overview of EBOV GP in complex with 11886 and 11883 Fabs. **(A) Schematic representation of EBOV GP.** Mucin-like domain (MLD), transmembrane domain (TM) and Cytoplasmic tail (CT) are deleted. Receptor binding region (RBR) and location of cathepsin cleavage loop (CL) are indicated in red and khaki respectively. The glycan cap (GC), GP2 N-terminus, internal fusion loop (IFL), heptad repeat 1 (HR1) and heptad repeat 2 (HR2) are colored blue, coral, purple, green and yellow respectively. GP1 (light grey) and GP2 monomers are linked by a disulphide bond. Hashed areas of schematic indicate areas of disorder in the structure. Signal peptide (SP) and membrane-proximal external region (MPER) are also indicated. **(B) The 3.0 Å cryo-EM map of the EBOV GPΔmuc ectodomain in complex with three 11883 Fabs (pink) and two 11886 Fabs (green) with no symmetry applied.** GP1 and GP2 are light and dark grey respectively. **(C) Surface representation of the top and side views of the model built into the map shown in (B)**. Fabs simultaneously bound to the GP trimer are numbered.

Consistent with the biochemical data and loss of binding to GP_CL_, 11883 contacts the top of GP1, making extensive contacts with the interior side of the GC (Asn238, Leu239, Thr269, Thr270, Gly271, Lys272, Leu273, Ile274, Glu287) via its CDRH1 and CDRH3 (**Figure 5Ai**). In addition, the CDRH3, CDRL1 and CDRL3 contact GP1 across the RBR (Glu112, Lys114, Pro116, Asp117, Gly118, Ser119, Ser142, Gly143, Thr144, Gln221) (**Figure 5Aii**). Among the contacts made on the GP1, Lys114, Gly118, Ser142, Gly143 and Thr144 are also residues used by GP to interact with NPC1 (**Figure 5A**). The 11883 epitope overlaps the footprint of mAb114 (an RBR-binding mAb that is approved for treatment of EVD ^7^) with shared contacts on the GP1 core and GC **(Figure S4A**). However, unlike 11883, mAb114 retains binding to EBOV GP_CL_ ^39^; this difference is explained by the additional contacts on the GC with 11883 compared to mAb114, and the greater GP1 core contacts with mAb114 compared to 11883. Hence, 11883 likely neutralizes by preventing NPC1 binding via competing for the RBR and/or by limiting GC removal.

**Figure 5.**
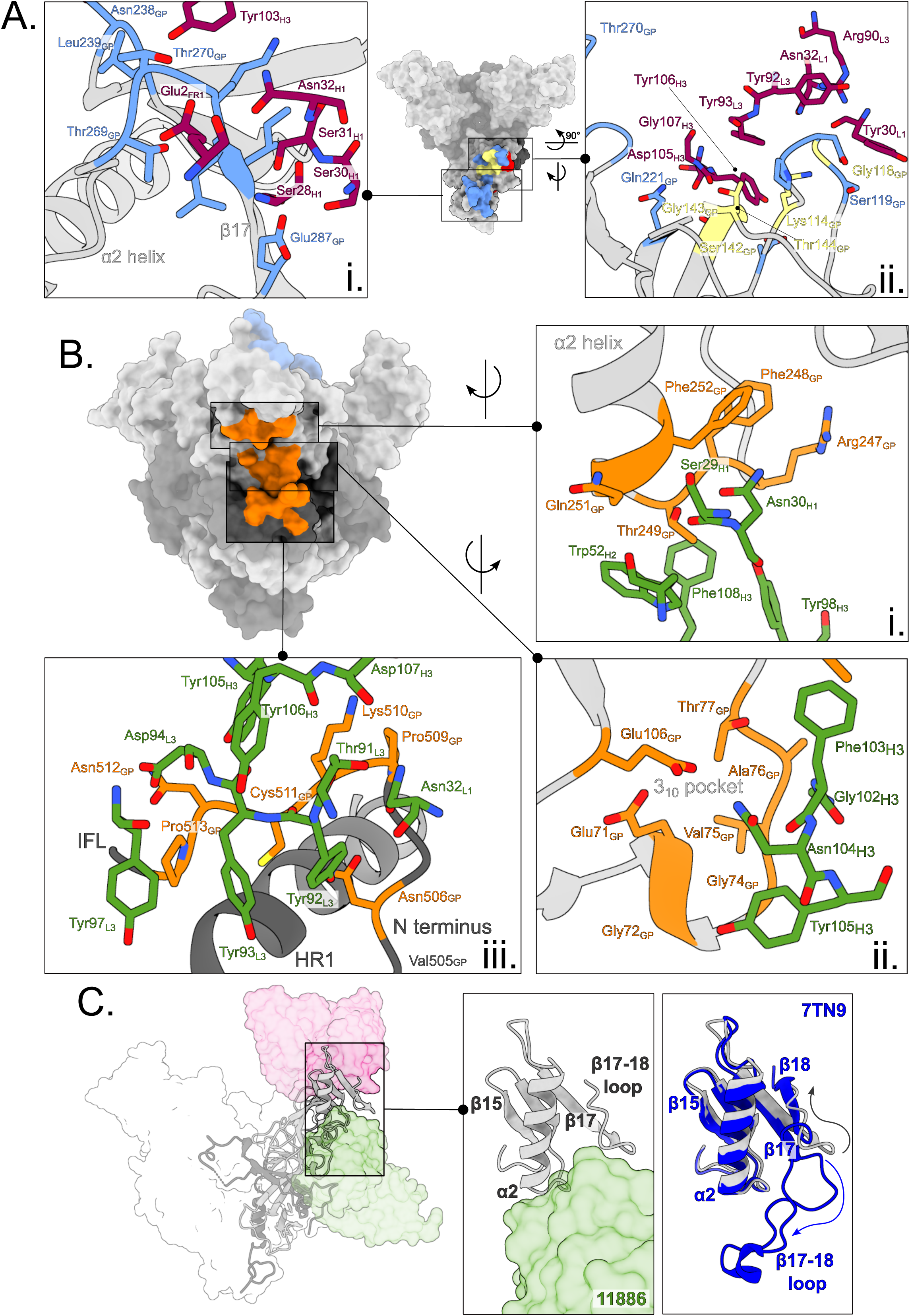
11883 binds across the receptor binding site and glycan cap, whereas 11886 binds across the glycan cap, 3_10_ pocket and GP2 N-terminus of a GP1/2 protomer. **(A) Surface representation of EBOV GPΔmuc trimer with 11883 epitope footprint highlighted.** 11883 contact residues on GP1 are colored blue and NPC1 receptor binding site residues are in red, with residues in common between the two footprints colored yellow. Where shown as sticks in insets, R groups of residues are coloured by heteroatom, with 11883 residues shown in dark pink. (i) Top view of GP showing extent of 11883 epitope footprint across the glycan cap (GC) and (ii) side view showing 11883 contacts in receptor binding site of GP1. **(B) Surface representation of EBOV GPΔmuc trimer with the tripartite 11886 epitope footprint highlighted in orange** (i) 11886 HC residues (green) interact with the base of the α2 helix and neighbouring residues of the GC. (ii) 11886 HC residues (green) interact with residues in the 3_10_ pocket of GP1. (iii) 11886 LC residues contact the GP2 N-terminus adjacent to the end of the IFL and above HR1. 11886 HC residues 130-132 make additional contacts with GP Lys510. **(C) 11886 displaces the β17-18 loop from the 3_10_ pocket.** A comparison of the position of 11886 Fab (green) and the start of the largely unresolved and missing β17-18 loop in this model (light grey) with that of the well-resolved β17-18 loop in the model of GP in complex with the Inmazeb mAb cocktail (dark blue) (PDB: 7TN9).

### 11886 recognizes a tripartite epitope, bridging the GC, the 310 pocket, and the GP2 N-terminus

Within this same model, mAb 11886 buries 1,030 Å^2^ of molecular surface on GP. First, consistent with a dependency on the presence of the glycan cap for binding to GP, 11886 CDRH3, CDRH2 and CDRH1 contact residues 247-249 and 251-252 at the base of the α2 helix and preceding β16-α2 loop in the GC (**Figure 5Bi**). Second, at the centre of its epitope, CDRH3 residues bind into the 3_10_ pocket of the GP (**Figure 5Bii**). A hydrogen bond is formed between Asn104_H3_ and E106_GP_ at the edge of the pocket. Multiple hydrophobic interactions are made directly with the 3_10_ helix itself (residues 71-75) between Tyr105_H3_ and Gly72_GP_ and Gly74_GP_, and between Gly102_H3_ and Val75-Ala76_GP_. The aromatic ring of Phe103_H3_ sits between the top of the 3_10_ pocket and the α2 helix forming interactions with both Thr77_GP_ and Thr249_GP_. Third, the CDRL1, CDRL3 and CDRH3 contact the GP2 N-terminal domain after the end of the IFL (**Figure 5Biii**). Here, the CDRH3 sits above the GP2 N-terminal domain and Tyr105_H3_ (which contacts the 3_10_ helix above) and also interacts with Asn512 and Lys510. The Lys510_GP_ side chain sits between the two antibody chains, allowing it to also interact with residues in the CDRL1, CDRL3 and CDRH3 (Asn32_L1_, Thr91_L3_, Tyr92_L3_, Tyr106_H3_ and Asp107_H3_). Hydrophobic and electrostatic interactions between Tyr97_L3_, Asp94_L3_, Tyr93_L3_ and Tyr92_L3_ with 513-511 of the GP, with an additional electrostatic interaction between Tyr92_L3_ and Asn506_GP_, complete the interface between the CDRL3 and the portion of the GP2 N-terminus sitting above HR1.

Notably, the lack of change in 11886 binding to chimeric GP with the MARV IFL (in which residues 512-555 of EBOV GP were replaced with the equivalent sequence from MARV GP) suggests that the interactions with Pro513 and Asn512 are either not necessary for 11886 binding to EBOV GP or are compensated for by interactions with the equivalent Ala513 and Asp512 residues in the chimeric GP. Residues Asn506, Lys510 and Cys511 were also maintained between the chimeric GP and EBOV GP and the extent of interactions with Lys510 may have compensated for changes further along the GP2 N-terminus.

Whilst sharing the 3_10_ pocket and contacts in the GP2 N terminus with previously described mAbs EBOV-520 and ADI-15946 ^22,28^, the footprint of 11886 sits further from the fusion loop and higher up the arête of the GP1,2 protomer with more extensive GC contacts (**Figure S4B**). In addition, 11886 appears to induce changes to the pre-fusion GP structure. In this map, the β17-18 loop is largely unresolved. Comparison with PDB:7TN9 ^26^, in which the β17-18 loop is resolved and occludes the 3_10_ pocket, shows that when 11886 is bound to the GP, the CDRH3 of the mAb directly clashes with the β17-18 loop position in 7TN9 (**Figure 5C**); these data suggest the β17-18 loop is not occluding the 3_10_ pocket in the 11886-bound structure. Further, Glu287 of the β17-18 loop is displaced ∼25 Å from its position in 7TN9, such that it swings to the top of GP and forms a contact with 11883 (**Figure 5Ai**). Furthermore, the tripartite epitope of 11886, linking the GP2 N-terminus to the 3_10_ pocket and GC of GP1 likely pins the GP in a pre-fusion conformation: neutralization therefore likely occurs via prevention of conformational changes required for membrane fusion.

In addition to structural investigation of the 11886 epitope, biologically contained ΔVP30 EBOV was passaged in the presence of mAb 11886 and viruses to select for mutations that allow escape from neutralization by the mAb. Escape viruses were sequenced and mutations V505I and T402I in the GP identified. Each mutation is individually sufficient to abrogate neutralization of virus by 10 μg of 11886 (**Figure S4C**). Whilst neither residue is defined as a contact residue on the GP for 11886 as determined by cryo-EM, the proximity of Val505 to the 11886 footprint in the GP N terminus, adjacent to contact residue Asn506, suggests that the substitution is likely to disrupt this portion of the 11886 footprint on the GP (**Figure 5Biii**). Interestingly, Val505 is not conserved across the GP species that 11886 is able to bind to and neutralize (**Figure S4D**), suggesting that the ability of this mutation to lead to escape from neutralization is determined by the change to the bulkier isoleucine, rather than the change from valine in EBOV GP. In contrast, the other escape mutation, T402I, is not defined in the cryo-EM structure as it is within the MLD, and is unlikely to contacted by 11886 due to the positioning of the 11886 footprint along the outer edge of the GP trimer and down to the base of the GP. It too is not conserved across GP species. This mutation may affect processing of the GP or positioning of the MLD, rather than directly interfering with 11886 contact residues.

In summary, 11883 binds the GC and RBR interior of the GP chalice, competing for NPC1 contact residues. 11886 binds down the side of the GP chalice making contacts with the GC, the GP1 3_10_ pocket (preventing its occlusion by the β17-18 loop) and the GP2 N terminus. The high degree of conservation between *Ebolavirus* species at these sites is consistent with the ability of these mAbs to recognise all GPs tested (**Figure S4D**).

### 11886 interacts synergistically with a range of anti-GC/RBR mAbs to increase their breadth of neutralization

Co-operative interactions between pairs of mAbs binding to the GP have previously been reported, in particular, where one partner is a cross-neutralizing mAb that binds across the 3_10_ pocket ^21,27,28^. Our previous data identified mAb 11886 as the most promising broadly-neutralizing clone, and show that it contacts the 3_10_ pocket. We therefore next evaluated the potential of this mAb to interact synergistically with other mAbs targeting non-overlapping epitopes around the GP in order to determine the potential value of including 11886 in a future multivalent mAb cocktail (**Figure 6 and S5**). Other test antibodies were selected based on lack of competitive binding to GP with 11886, breadth of binding to different *Ebolaviruses*, and range of neutralization potencies against the EBOV, SUDV and BDBV GP S-FLU pseudotype viruses. Partner mAbs were titrated alone (red lines) and in combination with a fixed concentration of 11886 (blue lines) in a series of GP pseudotyped S-FLU neutralization assays. A Bliss additivity model ^40^ was used to predict the neutralization of the mAb mixture if their effects were independent and additive (solid grey lines); neutralization above these predicted values suggests a synergistic interaction between the partner antibodies.

**Figure 6.**
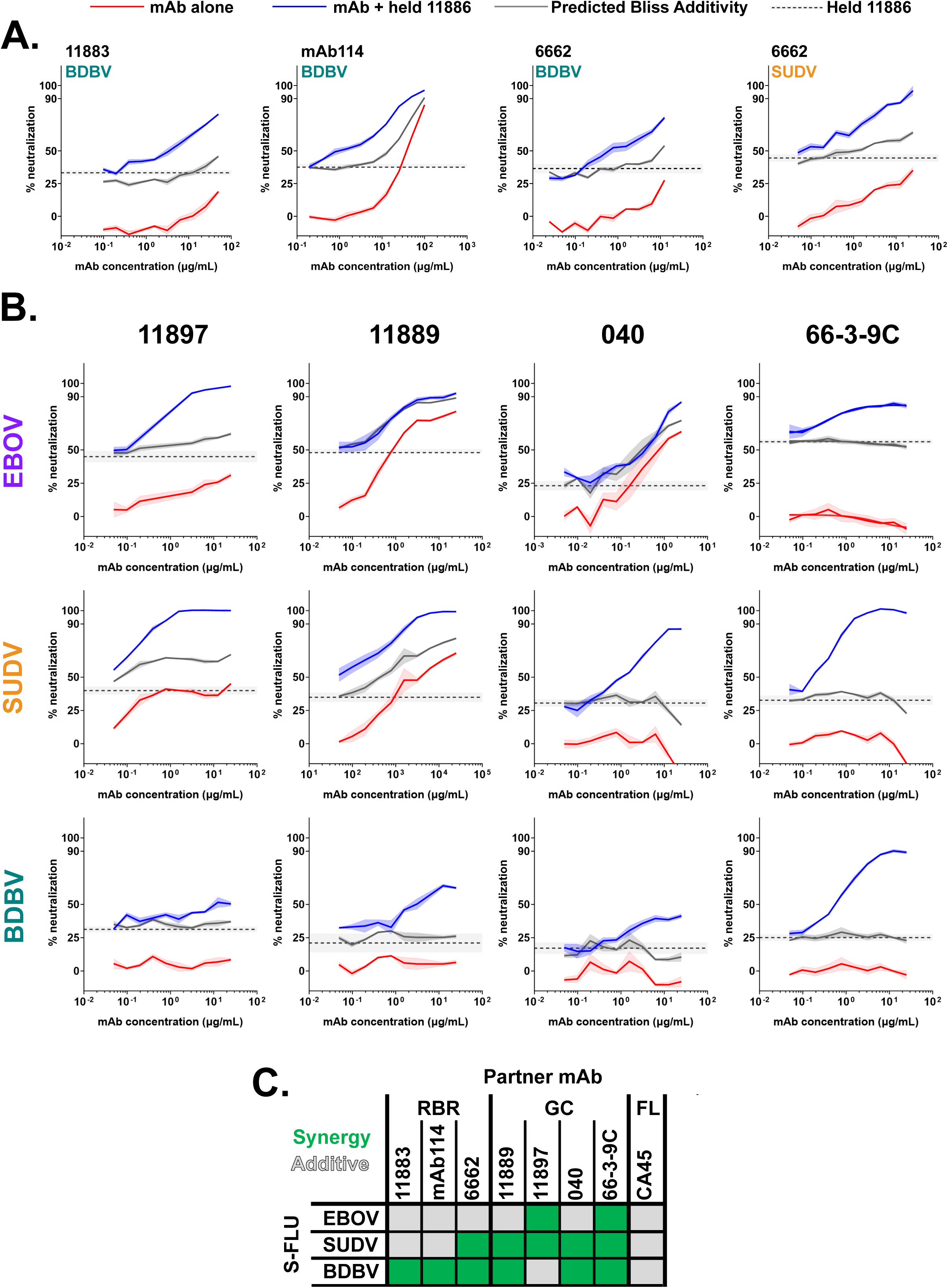
11886 potentiates a range of GC and RBR binding broadly reactive partner antibodies across Ebolavirus species in a pseudovirus neutralization assay. **(A) 11886 potentiates RBR mAbs 11883, mAb114 and 6662.** **(B) 11886 potentiates non-competing GC mAbs 11897, 11889, 040 and 66-3-9C.** For all data, a representative experiment of at least two independent repeats is shown. Dotted line represents the mean inhibition given by a held fixed concentration of 11886 with 95 % confidence limits shown in shaded grey. Solid lines represent mean of triplicate wells within an experiment, with shaded areas indicating the standard error. Red lines indicate the inhibition given by a titration series of the partner mAb alone. Blue lines indicate the inhibition given by a titration series of the partner mAb plus the held concentration of mAb 11886. Grey line represents the calculated Bliss Additivity value which assumes a purely independent and additive interaction between the partner mAb and 11886. Held concentration of 11886 ranged from 0.2-0.6 μg/mL between experiments to achieve target 20-45 % inhibition in a given assay. **(C) Summary of interactions between 11886 and partner mAbs tested against EBOV, SUDV and BDBV GP S-FLU pseudoviruses.** Green squares indicate a synergistic interaction between 11886 and mAb in neutralizing the virus, grey squares indicate independent and additive interactions.

Initially we observed synergy with anti-RBR mAbs starting with clone 11883. Alone this mAb could not neutralize BDBV GP S-FLU (<20 % neutralization at highest concentration tested), however, when partnered with a fixed concentration of 11886, a clear synergistic interaction occurred leading to neutralization up to ∼80 % (**Figure 6A**). Similar experiments with EBOV and SUDV GP S-FLU identified an additive, as opposed to synergistic interaction (**Figure S5A**). Next we tested mAb114, an approved monovalent therapy for EVD caused by EBOV, but likely with limited efficacy against other *Ebolavirus* species ^11^. Interestingly, antibodies 11883 and mAb114 share some of the same footprint in the RBR (**Figure S4A**) and notably the combination experiments of 11886 and mAb114 showed a series of very similar outcomes to those seen with 11883 (**Figure 6A, S5A**). Finally, mAb 6662 binds the RBR and is component of a mAb cocktail that protected guinea pigs from EBOV challenge, however, alone it is only able to partially neutralize the viruses tested here ^34^. In this case, 11886 and 6662 interacted synergistically against both BDBV and SUDV GP S-FLU, whilst showing additivity against EBOV GP S-FLU (**Figure 6A, S5A**). These data clearly identified that 11886 could increase the neutralization breadth and potency of mAbs that bind to the RBR when used in a bivalent combination.

Due the lack of competition between 11886 and IFL stem binding mAb CA45 ^31^, we next assessed the combination of 11886 and CA45 in this assay. Here we observed exclusively additive interactions when using the mAb combination (**Figure S5B**).

Finally, we tested whether 11886 can increase the neutralization breadth of mAbs that bind to the wider GC domain. The exact epitopes of 11897 and 11889 are unknown, but as shown in **Figures 2 and 3**, their footprints encompass the GC. Antibodies 040 and 66-3-9C are also components of the reported protective cocktail containing RBR mAb 6662, and bind to non-competing epitopes in the GC ^34^. In particular, 66-3-9C binds to the β17-β18 loop. All these mAbs are broadly reactive, but not broadly neutralizing when tested individually. However, when partnered in combination with 11886, all showed synergistic neutralization of at least two pseudoviruses, with the remaining combinations being additive (**Figures 6B, 6C**). Most notably, mAb 66-3-9C alone is non-neutralizing, but in combination with 11886 displayed 90-100 % synergistic neutralization of all three viruses (**Figure 6B, 6C**).

### Inclusion of 11886 and 11883 improves breadth of neutralization of cocktails of antibodies

The ability of *Ebolaviruses* to generate mutations allowing escape from neutralization by one (as demonstrated in **Figure S4C** for mAb 11886 in this study) or even cocktails of two broadly reactive GP targeting mAbs ^19,25^ implies that the development of therapeutics for EVD should focus on cocktails of 3 or more mAbs. Therefore, we next tested 3 or 4 component mixes of antibodies in the S-FLU pseudovirus assay (**Figures 7 and S6**). Each cocktail contained non-competing mAbs (combined in an equal ratio) against the GC/RBR, the GC β17-18 loop and the base of the GP. A cocktail of 11886 + 11883 + 66-3-9C (Mix 1) was selected on the following basis: (1) due to the breadth and potency of neutralization of 11886 and 11883; (2) the observed synergy of 11886 and 66-3-9C; and (3) the lack of competition between the three mAbs. The cocktail of 6662 + 040 + 66-3-9C + 6541 (Mix 2) contains cross-reactive mAbs and has previously been shown to be protective against EBOV challenge in guinea pigs ^34^, however it is unable to neutralize SUDV S-FLU *in vitro*. Cocktails of mAb114 + 66-3-9C + 11886 (Mix 3) and 6662 + 040 + 66-3-9C + 11886 (Mix 4) were also included for comparison to investigate the best combination of these component mAbs. All cocktails were prepared with equal amounts of each component mAb and the same total amount of antibody titrated in the assay (starting with 0.625 μg total antibody at 12.5 μg/mL). The cocktails were assessed in N=3 replicate experiments head-to-head with mAb 11886 alone. IC50 and IC80 values (concentration of antibody required to reach 50 % and 80 % neutralization of virus, respectively) were calculated from titration of the antibody mixes. For comparison, IC50 values from previous experiments using each of the other component mAbs alone were calculated and are shown (**Figures 7 and S6**).

**Figure 7.**
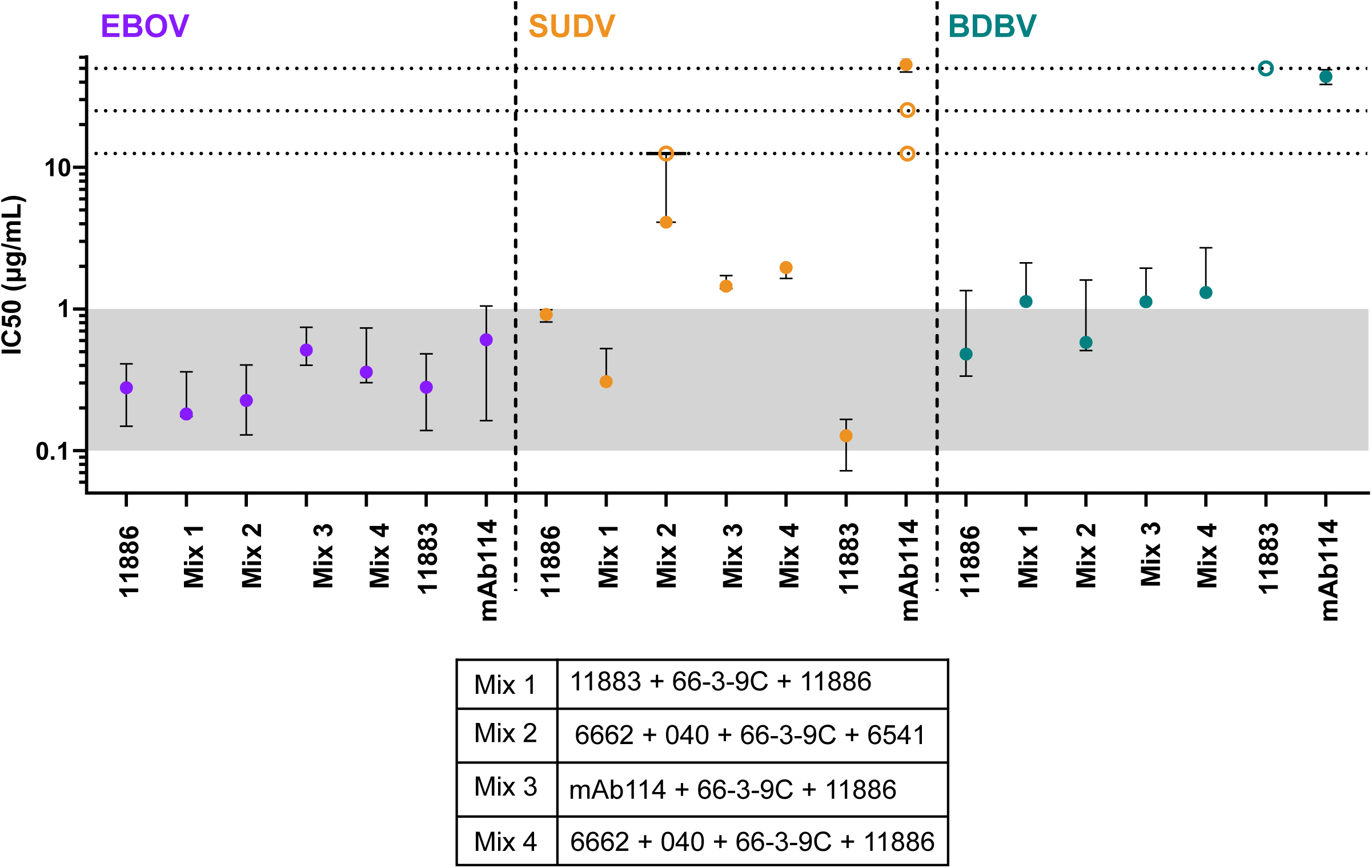
Cocktails of antibodies containing 11886 and 11883 have improved breadth of neutralization of pseudoviruses. Antibody cocktail mixes were tested for neutralization of EBOV (purple), SUDV (orange) and BDBV (teal) GP coated S-FLU pseudoviruses. Median and 95 % CI of the IC50 values (concentration of antibody required to achieve 50 % virus neutralization) for antibody mixes and 11886 alone were calculated from N = 3 experiments. For comparison, median and 95 % CI of IC50 values of individual antibodies 11883 and mAb114 calculated from N=2 to N=5 experiments run separately are also shown. Open symbols represent results where 50 % neutralization of virus was not achieved at the highest concentration of the test antibody assayed in that experiment. Horizontal dotted lines denote 12.5, 25 and 50 μg/mL of total antibody respectively. Shaded region demarcates IC50 concentration range 0.1-1 µg/mL.

All mixes neutralized EBOV S-FLU pseudovirus with similar IC50 and IC80 values to each other and to 11886, 11883 or mAb114 alone (95% CI intervals for all IC50 values are within ∼0.14-1.0 μg/mL). Similarly, all mixes neutralize BDBV S-FLU pseudovirus with similar IC50 and IC80 values to each other and to 11886 alone (IC50 values median and 95% CI range ∼0.3-2.7 μg/mL).

In addition, Mix 1 showed improvement over the other cocktails in neutralization of the SUDV S-FLU pseudovirus. As expected, Mix 2 was unable to neutralize SUDV S-FLU effectively at the concentrations tested, due to the inability of the component mAbs to neutralize 50% of virus individually. In contrast to the other cocktail component mAbs (including mAb114), 11883 and 11886 can both individually neutralize SUDV S-FLU. Therefore Mix 3 and Mix 4, which each contain one of 11883 or 11886 show IC50 values of SUDV S-FLU of ∼1.4-2 µg/mL. Further, Mix 1, which contains both 11886 and 11883 shows IC50 values of ∼0.3-0.5 µg/mL. Therefore, candidate cocktails including 11886 and 11883 show improved breadth of neutralization of GP-coated pseudoviruses compared to the approved monotherapy mAb114 and a previously published cocktail (Mix 2) ^34^.

## Discussion

Currently approved therapies for EVD consist of mAbs that target the EBOV GP, with likely limited efficacy against other species of *Ebolavirus*. These therapies, whilst effective in reducing overall mortality from EVD during an outbreak of EBOV, showed reduced efficacy in cases with delay to treatment or higher viral loads ^11^. In response, the field has continued to develop a new pipeline of more potent and broadly reactive mAbs that show efficacy against EBOV, SUDV and BDBV in animal models. These therapies could be tested for efficacy in the event of a future outbreak of EVD.

Here we isolated and characterized mAb 11886, a novel antibody that is broadly neutralizing and has the potential to improve the efficacy of other mAbs targeting the GP across the RBR and GC when combined in a cocktail, thereby increasing both potency and breadth of action. The 3.0 Å cryo-EM structure we present here delineates the tripartite epitope of 11886 down the arête of the GP chalice, bridging and pinning together the GP2 N-terminus, GP1 core and the GC into a pre-fusion conformation, thereby likely preventing the rearrangements required for receptor binding, membrane fusion and infection. This epitope is distinct from previously described broadly neutralizing antibodies that also bind the 3_10_ pocket and GP2 N-terminus, and may enhance neutralization by a range of GC and RBR mAbs by holding the GC in a preferable conformation.

Other broadly neutralizing mAbs that bind the conserved sites of vulnerability in the GP N-terminus and 3_10_ pocket of the GP1 core have been described, such as ADI-15946 (and its improved derivative ADI-23774^AF^) ^28,41^ and EBOV-520 ^21,22^. However, these mAbs, unlike 11886, do not interact with the GC in a significant manner. Whereas 11886 loses binding to GP_CL_, ADI-15946 and EBOV-520 recognize GP_CL_; both mAbs maintain (or improve) neutralization of viruses with pre-cleaved GP ^21,22,28^, whereas in this study, 11886 lost neutralization of pseudotyped viruses pre-treated with THL. This dependency on the presence of the GC for 11886 binding is explained by the interaction between 11886 heavy chain and the α2 helix and β16-α2 loop.

The 3_10_ pocket is a region of the GP between the GC and GP2 that is usually occupied by the β17-18 loop (residues 287-291) of the GC. 11886, ADI-15946 and EBOV-520 all interact with a key conserved E106 residue that sits to the top left of the pocket above the 3_10_ helix that forms the majority of the pocket (residues 71-75) ^21,28^. Binding to the 3_10_ helix, holding it in the pre-fusion conformation and preventing the rearrangement of these residues that occurs after GC cleavage and receptor binding, may contribute to neutralization by these mAbs. 11886, like ADI-15946 and EBOV-520, protrudes into the pocket resulting in displacement of the β17-18 loop. Various mechanisms that enhance displacement of the loop from the pocket also enhance binding of EBOV-520 and ADI-15946; these include remodelling by partner mAbs, introduction of mutations into the loop to reduce interaction with the 3_10_ pocket, and complete removal of the loop ^21,28^. As 11886 requires the GC to be present to bind to the GP, mechanisms that displace the loop, but retain the normal positioning of the α2 helix and β16-α2 loop could also enhance 11886 access to the 3_10_ pocket and binding to GP. FVM09, a non-neutralizing mAb that binds to the β17-18 loop enhances activity of ADI-15946 by displacing the loop ^27,28^. 66-3-9C binds to the same GEWAF (residues 286-290) peptide as FVM09, and similarly is non-neutralizing alone ^32,34^. This comparison led us to test if 66-3-9C and 11886 also interact synergistically analogous to FVM09 and ADI-15946. Clear synergy between 66-3-9C and 11886 was observed across all viral species tested. At least part of this synergy may therefore be due to similar co-operative effects resulting in enhanced displacement of the loop from the pocket, thereby resulting in enhanced access to the loop residues for 66-3-9C and/or the 3_10_ pocket residues for 11886. Nevertheless, even without 66-3-9C present, 11886 is able to mediate significant rearrangement of the position of loop. In the cryo-EM structure of 11886 and 11883 in complex with GP, the displacement of the β17-18 loop by 11886 is so complete that residue 287 (in the centre of the GEWAF peptide), which usually sits below the GC alongside the 3_10_ pocket, is able to interact with FR1 of 11883 binding at the top of the GC. The co-operativity of EBOV-520 and GC mAb EBOV-548 involves remodelling of the GC to enhance EBOV-520-mediated displacement of the β17-18 loop improving its binding to the EBOV GP 3_10_ pocket; indeed, a structure of the two antibody Fabs in complex with GP shows upwards displacement of the GC α2 helix and β17 strand compared to unliganded structures ^21^. In contrast, in the structure presented here with 11886 and 11883 bound to the GP, the GC more closely resembles the unliganded structure or the mAb114 bound GP in regions where the GC is well-resolved. This suggests a different mechanism in which the EBOV GC is held in standard pre-fusion configuration by 11886 except for the displacement of the β17-18 loop.

Overall comparison of the footprints of these mAbs illustrates that 11886 binds further up the side of the GP chalice and across the GP away from the base of the IFL of the same protomer as compared to both ADI-15946 and EBOV-520. However, it still makes several contacts with the GP2 N-terminus. The dependency of 11886 binding on the presence of the GC, but lack of change in binding when residues 512-555 are changed (the opposite to ADI-15946 and EBOV-520), illustrates that despite their overlapping footprints in the 3_10_ pocket and GP N-terminus, these mAbs have distinct modes of binding to the GP.

Cocktails of antibodies provide opportunities not only for reduced risk of viral escape and increased breadth of possible Fc effector function recruitment, but also for co-operativity to enhance the potency of the component antibodies. 11886 interacts synergistically with a range of mAbs with different epitopes across both the RBR and GC. This suggests there is a general mechanism whereby 11886 is potentiating multiple different antibodies that bind across this broad region of the GP. Co-operativity has been shown with other mAb pairings against EBOV GP, for example EBOV-520 and EBOV-548, and ADI-15946 and FVM09 ^21,28^. However, both of these examples involve specific remodelling of the interaction between the β17-18 loop and the 3_10_ pocket. Here we show that enhanced neutralization can occur when 11886 is partnered with a range of different mAbs that do and do not interact with the β17-18 loop with epitopes across the GC and RBR region. We therefore posit that the pinning of the bottom of the GC to the GP1 core and GP2 N-terminus by 11886 likely holds the GC, and by extension the MLD, in a conformation that enhances binding to the top of the GC and RBR. 66-3-9C has an epitope that includes residues in the β17-18 loop. As discussed above, the interaction between this mAb and 11886 may be distinct from the mechanism of potentiation of other antibodies that access the inner chalice of the GC and the RBR crest and have higher involvement of the β17-18 loop rearrangements. However, 66-3-9C also loses binding to GP when the MLD is removed ^34^. It may be that aside from the effects on the loop directly, any pinning of the GP by 11886 orders the wider 66-3-9C epitope including the MLD component.

In this study, more mAbs are enhanced against SUDV and especially BDBV, than are enhanced against EBOV. Many of the partner antibodies tested in this study are individually more neutralizing against EBOV than they are SUDV or BDBV. Therefore, this improvement in BDBV and SUDV neutralization when partnered with 11886 may simply reflect the difficulty in showing improvement of an already neutralizing mAb in the assay, i.e. it is easier to show enhancement when one antibody is poorly neutralizing alone. An alternative explanation, however, is that mAbs that bind and neutralize EBOV well, may struggle to neutralize BDBV and SUDV if there are differences in the arrangement of the GC and MLD in these GP species that restrict access to their epitopes. If 11886 is able to pin the BDBV and SUDV GPs in a more EBOV-like configuration, this may improve access to the epitopes of these antibodies and improve their neutralization of these species too.

This has implications for future therapeutic cocktail design. There is a clear need for more potent therapies against EBOV and for therapies with efficacy against other known, and yet to emerge, *Ebolavirus* species. We have shown that inclusion of 11886 and 11883 in cocktails of mAbs can improve the potency and breadth of action *in vitro* of current EVD therapies that include RBR or GC targeting mAbs such as mAb114. The difficulties of testing new therapies in the context of an outbreak of EVD, and the future ethical considerations surrounding offering experimental treatments versus now approved and tested therapies needs careful consideration. The addition of a single antibody to enhance the action of many may allow strategic supplement of more clinically developed therapies for assessment during outbreaks. As 11886 does not compete mAbs that bind the stem or tip of the IFL, it could be included in cocktails with mAbs targeting these epitopes. There are multiple cocktails under pre-clinical assessment including one IFL and one RBR mAb (FVM04/CA45 and 1C3/1C11 ^23,35^) that could warrant assessment of the addition of a 11886-like component.

## STAR Methods

## Resource Availability

### Lead contact

Further information and requests for resources and reagents should be directed to and will be fulfilled by the lead contact, Simon Draper.

### Materials availability

### Data and Code Availability

Structural data have been deposited at PDB and are publicly available as of the date of publication. Accession numbers are listed in the key resources table. All other data reported in this paper will be shared by the lead contact upon request. This paper does not report original code. Any additional information required to reanalyze the data reported in this paper is available from the lead contact upon request.

## Experimental model and study participant details

In this study, 12-16 week old, female, New Zealand white rabbits were housed in floor pens with temperatures in the room maintained between 16 and 20 °C with the same humidity with 12 h light 12 h dark cycled on the system from 7 to 7. Approval for use of animals for immunization was provided through the UCB Pharma, UK Animal Welfare and Ethical Review Body (AWERB) and the license was granted by the UK Home Office.

## Method details

### Rabbit immunizations

RAB-9 cells were cultured to a density of approximately 1 x 10^8^ cells/ 5-stack CellSTACK® Culture Chambers (Corning, 3319) in RAB-9 culture media. Cells were lifted using StemPro™ Accutase™ Cell Dissociation Reagent (Gibco, A1110501) and resuspended at 5 x 10^7^ cells/mL in Earle’s Balanced Salts (Sigma, E3024). Cells were incubated with plasmids containing full-length *Ebolavirus* GP sequences (EBOV GP (NP_066246.1), SUDV GP (YP_138523.1), BDBV GP (YP_003815435.1) or TAFV GP (YP_003815426.1)) and electroporated before seeding into flasks and incubated at 37 °C, 5 % CO_2_. Cells were lifted the next day and expression of antigens was confirmed by FACS via staining with human antigen-specific antibody and goat anti-human Fc-specific AffiniPure F(ab’)₂ Fragment Alexa Fluor 647 conjugate (Jackson 109-606-170). Fluorescence was measured using BD FACS Canto^TM^ 11 and analyzed with FlowJo3 software. A female New Zealand white rabbit 12-16 weeks old was immunized four times subcutaneously at two week intervals with a total mixed immunogen dose of 1 x 10^7^ RAB-9 cells, consisting of 2.5×10^6^ cells transfected with each of EBOV, SUDV, BDBV and TAFV GP. First dose only was adjuvanted with complete Freund’s adjuvant delivered at a separate site. Serum was sampled on each day of vaccination and monitored for binding to GP before the rabbit was euthanized by intravenous overdose of anaesthetic, and PBMC, spleen, bone marrow and lymph nodes harvested, 2 weeks post-final vaccination.

### Monoclonal antibody recovery

B cell culture and primary screening was conducted using UCB’s single-B cell platform, as described previously ^42^. Briefly, splenocytes were co-cultured with irradiated mutant EL-4 murine thymoma feeder cell line in B cell media with proprietary supplement, similar to culture as described by Lightwood *et al.* ^43^. Cell culture supernatants were screened for reactivity to *Ebolavirus* GPs expressed on both MDCK SIAT-1 cells and Expi293 HEK cells. Individual antigen-specific IgG secreting B cells were recovered using the fluorescent foci method (US Patent 7993864/ Europe EP1570267B1) using beads immobilized with recombinant purified GP ^44^. V region sequences were recovered by nested RT-PCRs, then subcloned into rabbit IgG and Fab expression vectors.

### Immunofluorescence assays (IFA) using GP expressing MDCK-SIAT 1 cell lines

MDCK SIAT-1 cells were seeded in 96 well flat, black skirted, tissue culture plates (Corning, 3904) at a density of 3×10^5^ cells per well in 100 µL D10 media and recovered overnight at 37 °C, 5 % CO_2_. Cells were washed in PBS to remove D10 media, and incubated in 50 µL mAb diluted in PBS for 1 h, 4 °C. Cells were washed then incubated in 50 µL 5 µg/mL anti-rabbit IgG Alexa Fluor 647 conjugate (Invitrogen, A21244) for 1 h, 4 °C. Plates were washed and 100 µL PBS added per well. If plates could not be read immediately, cells were fixed with 1 % formalin/PBS. Fluorescence was measured using a Clariostar plate reader (BMG Labtech) as previously described ^29^. R5.034 (a non-anti-GP mAb) ^45^ and 66-3-9C ^34^ were included as negative and positive controls respectively.

#### Antibody competition IFA

Competition experiments were conducted using an adapted version of the IFA. Biotinylated mAb1 at 5 µg/mL in PBS and unbiotinylated mAb2 at 50 µg/mL in PBS were mixed in equal volume prior to addition to cells. Each assay plate included competition controls where mAb1=mAb2 and additionally where mAb2 was a mAb against an irrelevant antigen or PBS only as a minimum competition control. Cells were washed then incubated in 50 µL 2 µg/mL Streptavidin Alexa Fluor 647 conjugate (Invitrogen, S21374) for 1 h, 4 °C. After washing, fluorescence was measured and the degree of competition was determined by: (X–Minimum binding)/(Maximum binding – Minimum binding), where ‘X’ is binding of the biotinylated mAb in presence of competing mAb, ‘minimum binding’ is the signal from the biotinylated mAb in the presence of self (unbiotinylated) and ‘maximum binding’ is the signal from the biotinylated mAb in presence of a non-competing non-GP mAb.

#### IFA with thermolysin digested-GP

Binding to THL digested GP was determined using an adapted version of the IFA. After seeding and recovery, cells were washed and incubated in 100 µL 0.25 mg/mL THL (Sigma, P1512) in HM buffer (20mM MES, 20mM HEPES, 130mM NaCl, 2mM CaCl_2_, pH7.5) or HM buffer alone for 1 h, with gentle shaking. After this and every subsequent incubation, cells were washed with plates spun at 500 x *g* and supernatant aspirated between washes. Cells were incubated with 50 μL 10 µg/mL mAb or 10 μg/mL biotinylated NPC1 protein in PBS, and incubated for 1 h at room temperature (RT), then anti-rabbit IgG Alexa Fluor 647 conjugate (Invitrogen, A21244) or anti-human IgG Alexa Fluor 647 conjugate (Invitrogen, A21445) or Streptavidin Alexa Fluor 647 conjugate (Invitrogen, S21374), and wheat germ agglutinin (WGA) Alexa Fluor 488 conjugate (Invitrogen, W11261) were added and cells incubated for 1 h, at RT with gentle shaking, protected from light. Plates were centrifuged before fluorescence read at both 625-30/680-30 nm and 488-14/535-30 nm. Gain was adjusted to give a ratio of approximately 1 in wells containing cells that had been stained with both WGA-488 and anti-rabbit IgG-647 conjugates only (i.e. no primary antibody added). Wells with too few cells as determined by WGA-488 signal were excluded from analysis. For inhibition of THL digestion assays, cells were pre-incubated with 50 μL 10 µg/mL mAb before incubation with THL, rather than after digestion.

### Immunoprecipitation of thermolysin digested GP expressed on cells

MDCK SIAT-1 cells expressing GP were seeded at 8-12 x10^5^ cells per well in 6 well tissue culture plates (Corning, CLS3516) in 2 mL D10 and incubated overnight, 37 °C, 5 % CO_2_. Next day, cell surface proteins were biotinylated using EZ-Link Sulfo-NHS-Biotin (Thermo Scientific, 21217). Media was aspirated from wells and cells rinsed with PBS. 0.5 mL 0.5 mg/mL biotin reagent was added per well and incubated at RT for 30 min. Biotin reagent was removed, and free biotin quenched by addition of 100 mM glycine. Glycine was removed and 0.5 mL THL diluted in HM buffer or HM buffer alone added per well. Plates were incubated with gentle shaking for 1 h at RT. THL was gently aspirated, replaced with lysis buffer (Alfa-Aesar, J62805.AK) containing protease inhibitors (Sigma-Aldrich, P8340), and plates incubated at RT with gentle shaking for 30 min. Cells were pelleted and supernatant incubated with 7.5 µg precipitating antibody and 100 μL Protein A Sepharose slurry (Sigma-Aldrich, P3391) (5% w/v in lysis buffer). Samples were incubated with rotation for 1 h at 4 °C. Samples were then centrifuged in a pre-cooled microfuge for 5 min at 13,000 rpm and 4°C, and supernatant discarded. The sepharose A pellet was then washed twice with 0.5 mL chilled lysis buffer, pelleted and supernatant discarded. Protein was eluted from Sepharose A by addition of 35 μL sample loading buffer (Sigma-Aldrich, S3401-10) and incubation at 80 °C for 5 min. Samples were then centrifuged and supernatant loaded into Bolt 4-12% Bis-Tris Plus gel (Invitrogen). Electrophoresis was conducted in MES buffer at 100-120 V for 40 min with protein standards (NEB, P7119S, or Bio-Rad, #1610373). Proteins were transferred to nitrocellulose membrane and membranes blocked in 5% milk powder/PBS overnight at 4 °C. Membranes were washed three times with TBS (incubated 5-10 min per wash), then incubated in a 1 μg/mL dilution of Streptavidin Alexa Fluor 647 (Invitrogen, S21374) for 1 h at RT with rocking. Membranes were washed three times with TBS (incubated 5-10 min per wash) then imaged using an iBright FL100 (Thermo Fisher Scientific) with the auto exposure setting.

### S-FLU virus microneutralization assay

In 96 well, black-skirted, flat bottom tissue culture plates, 50 µL S-FLU viruses pseudotyped with *Ebolavirus* GP (as described in Xiao *et al.* ^29^) in Viral Growth Media (VGM) were incubated with 50 µL mAb diluted in PBS for 2 h at 37 °C and 5 % CO_2_, before addition of 100 µL MDCK SIAT-1 cells in VGM (3 x 10^5^ cells/mL). For assessment of cocktail mixes, antibodies were pre-mixed in equal ratio prior to addition of virus. Virus was used at a dilution previously determined to give maximum infection of cells. After 20-24 h incubation at 37 °C and 5 % CO_2,_ supernatant was aspirated from cells, and cells fixed with 100 µL 10% formalin for 30 min at 4 °C. Formalin was removed and 100 µL PBS added to each well. Fluorescence (GFP or mCherry) was read using a Clariostar plate reader (BMG Labtech) as described in Rijal *et al.* ^34^. Maximum infection was determined from cells infected with viruses pre-incubated with PBS only or a non-GP binding antibody. Minimum signal was determined from uninfected cells. Curves were fitted using GraphPad Prism 9.3.1 using a four-parameter nonlinear regression [Inhibitor] vs. response - Variable slope model. IC50 and IC80 values were determined by interpolation.

#### Thermolysin cleaved S-FLU virus assay

For the thermolysin cleaved virus neutralization assay, stocks of virus were divided into two and incubated with or without 250 µg/mL THL in VGM for 1 h at RT. Viruses were then buffer exchanged into fresh VGM using 100 kDa MWCO spin filter (Amicon, UFC910008) and resuspended in the same volume of VGM before addition to plates and incubation with mAbs. Background fluorescence from uninfected cell controls was subtracted and change in fluorescence calculated by subtracting fluorescence obtained when mAbs were incubated with undigested virus from fluorescence when mAbs were incubated with digested virus.

#### Synergy S-FLU virus assay

To determine interactions between pairs of mAbs, the above experimental set up was adapted as follows. Per assay plate, mAb1 was titrated in triplicate alone and in the presence of held mAb2. mAb2 was held at a concentration selected to give a target neutralization of 20-40 %. Bliss Additivity was calculated using values determined by mAb1 only and mAb2 only wells (at least 8 replicates per plate) ^40^. Calculated additivity was then compared to the experimentally determined neutralization by mAb1+mAb2 to determine if mAb pairings were additive or synergistic.

### Wild-type neutralization assay

Serial dilutions of mAb were incubated with 100 TCID_50_ of either EBOV Mayinga (GenBank accession: NC_002549, 8A phenotype), EBOV Makona (GenBank accession: KJ660347) or SUDV Boniface (GenBank accession: FJ968794), before Vero E6 cells (ATCC Cat. No: CRL-1587, passage 52) were infected with the mAb-virus mixture. Formation of cytopathic effect (CPE) was monitored after 7 days (Mayinga and Sudan), or 9 days (Makona) and compared with the CPE in untreated infected cells. The ovine polyclonal antibody EBOTAb (^46^ MicroPharm Ltd, Batch: 170315)) served as control. Virus neutralization titers (VNT) were calculated as geometric mean titres (GMT) of the reciprocal value of the last serum dilution at which inhibition of the CPE on infected Vero E6 cells was detectable, in comparison to the virus control. The initial concentration was 12.5 µg/mL and was therefore the upper limit of detection (ULOD). Lowest concentration tested was 0.006 µg/mL and was therefore the lower limit of detection. Assays were conducted under BSL4 conditions at the Philipps University Marburg.

### Escape mutant generation

EbolaΔVP30 viruses (biologically contained Ebola viruses in which the VP30 open reading frame has been replaced with a green fluorescent protein (GFP) reporter gene) were generated and propagated as previously described ^47^. To generate viral escape mutants, EbolaΔVP30 virus was propagated three consecutive times for 6 days in reduced medium (2% FBS, 1x MEM with antibiotics and supplements) with increasing concentrations of mAb 11886 (0.63, 2.5, and 5.0 µg/mL, respectively). Individual resistant plaques were selected, viruses isolated and virus stocks generated in the presence of 10 μg/mL mAb. Viral RNA was recovered from cell culture supernatant of virus stock (RNeasy minikit; Qiagen 74106), and viral GP gene amplified by RT-PCR (Verso 1-Step RT-PCR kit; Thermo Fisher Scientific AB-1455), cloned and sequenced as previously described ^48^.

### Protein Expression and purification

#### Expression and purification of mAbs and Fabs

Recombinant rabbit mAbs and Fabs were expressed in HEK293F cells using ExpiFectamine™ 293 Transfection reagent (Gibco, A14525). Human mAbs were expressed in ExpiCHO-S™ cells using an ExpiCHO™ Expression System Kit (Gibco, A29133). IgG were purified from cell culture supernatant 6-7 days after transfection via Protein A affinity chromatography using 2 mL or 5 mL column HiTrap MabSelect SuRe Protein A column (GE Healthcare, Life Sciences) dependent on culture volume. Columns were washed and IgG were eluted in 0.1 M sodium citrate, pH 3.4. Eluted fractions were neutralized with 2 M Tris/HCl pH 8.0. R5.034 and R5.014 were produced as described previously ^45,49^.

Fabs were purified from cell culture supernatant 7 days after transfection using serial affinity chromatography. Fabs were initially purified using a HisTrap Excel 5 mL column (GE Healthcare, Life Sciences), and eluted in 5 mL 0.5 M NaCl / 10 mM PBS/ 250 mM Imidazole pH7.4; then further purified using Gammabind Plus Sepharose Protein G 25 mL column (GE Healthcare, Life Sciences), with 20 min contact time, and eluted in 0.1 M glycine, pH 2.7. Eluted Fab was neutralized using Tris HCl pH 8.5. Small volumes of pooled eluted IgG or Fab were buffer exchanged into PBS pH 7.4 and concentrated using Amicon Ultra Spin columns with a 30 kDa cut off membrane (Millipore, UFC905008) and centrifugation at 4000 x *g*, before sterile filtering. Eluted samples with high concentration or in excess of 10-20 mL volume were buffer exchanged using two HiPrep 26/10 Desalting columns (GE Healthcare, Life Sciences) in series, flow rate 10 mL/min. Concentration was measured by A280 (Nanodrop spectrophotometer) and High Performance Liquid Chromatography. Monomer purity was assessed by size exclusion chromatography (SEC) on an Acquity UPLC system with a BEH200, 1.7 μM, 4.6 mm X 300 mm column (Waters, 176003905) and developed with an isocratic gradient of 0.2 M phosphate, pH 7.0 at 0.3 mL/min.

#### Recombinant GP expression

For production of soluble GP used in B cell recovery and foci picking HEK293 cells were transiently transfected. For SUDV GP expression, HEK293F cells were transfected with a plasmid encoding SUDV GPΔTM (1-649, Sudan virus/H.sapiens-tc/UGA/2000/Gulu-808892) with C-terminal C-tag (EPEA) peptide ^50^. After 5 days, supernatant was harvested, clarified by centrifugation, and filtered. For EBOV GP expression, HEK293E cells were transfected with a plasmid encoding EBOV GPΔTM (1-649 H.sapiens-wt/GIN/2014/Makona-Gueckedou-C07) with C-terminal C-tag peptide. After 3 days, supernatant was harvested clarified by centrifugation and filtered. Supernatant was concentrated 10 x using tangential flow filtration and immediately purified. GP proteins were purified via C-tag affinity chromatography, using a C-tag XL affinity resin column (Thermo Fisher Scientific) and eluted in 2M MgCl_2_, 20 mM Tris, pH 7.4. Eluate was further purified by SEC in TBS using a HiLoad 16/600 200pg column (GE Healthcare, Life Sciences).

For structural work, a *Drosophila* S2 cell stable cell line expressing the EBOV GP ectodomain with a double Strep-tag at the C-terminus (EBOV Mayinga, GenBank Ascension: AAN37507.1, Residues 33-651) was generated and adapted for larger scale expression as described previously ^26^. At a cell density of 1 million cells/mL, secreted GP was induced with 0.5 mM CuSO_4_ and supernatant harvested after 4 days. BioLock reagent (Iba, 2-0205-050) was added to supernatant (1:400) and GP affinity purified using a StrepTrap HP column (GE, 28907548). Eluate was concentrated and further purified via SEC on a Superdex200 (GE, 28907548) column in 50 mM Tris pH 7.5, 150 mM NaCl.

### Structural methods

#### Complex generation

Purified GP at a concentration of 1.9 µg/µL was complexed with 3x molar ratio excess of Fab 11886 for 4 h at RT. The complex was purified on a Superdex200 (GE, 28907548) column in 50 mM Tris pH 7.5, 150 mM NaCl. Fractions containing the complex were pooled and concentrated to 2 µg/µL. To this, 5x molar ratio excess of 11883 Fab was added and incubated overnight at RT. The resultant zGP-11886-11883 complex was used directly for structural studies without further purification.

#### Cryo EM sample preparation and data collection

The zGP-11886-11883 complex was diluted to A280 absorbance value 1 for the purpose of freezing grids. 3 µL complex was mixed with 1 µL 20 µM lauryl maltose neopentyl glycol (LMNG) and immediately applied to glow discharged C-Flat™-2/1-3Cu-T50 (Electron Microscopy Sciences) holey grids at 4 °C and 100 % humidity inside a Vitrobot Mark IV. Grids were blotted for 7 s with a blot force of 0, and plunge-frozen in liquid ethane cooled by liquid nitrogen.

Frozen grids were imaged on a Thermo Fisher/Scientific Titan Krios (G3) equipped with a K3 direct electron detector and a BioQuantum energy filter (Gatan). 9552 movies were recorded at a magnification of 75000x and a total dose of 50 e/A2 using EPU software.

#### Cryo-EM map calculation, structure determination and structure refinement

CryoSPARC ^51^ was used to process all the data. CTF estimations were performed on motion corrected and gain subtracted micrographs. A blob picker employing a wide size range (120 Å-280 Å) was used on an initial subset of 1000 micrographs to pick particles. Several careful and stringent 2D classifications jobs were used to reduce the total number of particles to 30,391 which were fed into Topaz ^52^, a neural network based particle picker. Two *ab initio* 3D volumes were also generated with subsets of these particles. Using the Topaz train model, particles were then extracted at a box size of 512 from the whole dataset and then binned to a box size of 128 for faster computation. Post-extraction and binning, extensive 2D classification jobs were performed on these particles to obtain a final set of 120,051 particles out of which 30,000 particles were used to generate two additional *ab initio* volumes. Heterogeneous refinements using the 120,051 particle set were performed on the four *ab initio* volumes among which one showed promising features for most of the bound Fab fragments. 66,550 particles were classified into this volume during heterogeneous refinement. This final set of 66,550 particles was re-extracted at a box size of 512 for final refinements. Attempts to refine without applying any symmetry showed the most promising features. Hence, refinements using C3 symmetry were abandoned. The final 3 Å map was generated using Non-uniform refinements after Local and Global CTF refinements were performed.

An initial model was generated using the PDB ID: 7TN9. This was docked into the map obtained from cryoSPARC. Several rounds of Phenix ^53^ refinement with the appropriate sequences, following by manual building in Coot ^54^ resulted in the final structure. This was visualized using UCSF ChimeraX ^55^.

## Author contributions

Conceptualization: FRD, SJD, DJL, ART

Data curation:

Formal Analysis: FRD, VR, HMK, AB, CH

Funding acquisition: SJD, DJL, ART, EOS, TS, PH

Investigation: FRD, VR, CH, HMK, TS, AB, AP, SH, KCLS, DP, RDA

Methodology: FRD, PR, LS, VOD

Project administration: FRD, SJD

Resources: FRD, VOD, PR, LS

Software: n/a

Supervision:

Validation: KH

Visualization: FRD, VR

Writing – original draft: FRD

Writing – review & editing: FRD, SJD, VR, EOS, ART, DJL, PR

## Acknowledgements

The authors are grateful for the assistance of Julie Furze, Penelope Lane, Fay Nugent, Jenny Bryant, Lana Strmecki, Sean Elias, Jing Jin, Geneviève Labbé, Daniel Alanine, Jordan Barrett, Kirsty McHugh and Julie Xiao, and to Sally Pelling-Deeves for arranging contracts (University of Oxford). The authors would also like to thank Dr. Verena Krähling (Institute of Virology, Philipps University Marburg) for the supply of wild-type *Ebolavirus* stocks used in this study and Gotthard Ludwig and Sebastian Schmidt from the BSL4 facility at the Philipps-University of Marburg for technical support. We also thank Sarfaraj Topia for support in protein production and Rebecca Munro for supporting the immunization work (UCB Pharma).

FRD held a UK Medical Research Council (MRC) iCASE PhD studentship [MR/N01796X/1]. SJD is a Jenner Investigator and held a Wellcome Trust Senior Fellowship [106917/Z/15/Z]. T.S. was supported by the German Research Foundation [197785619/SFB1021]. PR and AT were funded by the Chinese Academy of Medical Sciences (CAMS) Innovation Fund for Medical Science (CIFMS), China (grant no. 2018-I2M-2-002). EOS acknowledges NIH grant U19 AI109762.

## Declaration of Interests

DJL, VO’D are employees of UCB Pharma.

DJL holds stock and/or stock options in UCB Pharma.

FRD, SJD, DJL, VO’D, ART, PR and LS are inventors on patent applications relating to anti-*Ebolavirus*

GP antibodies.

