## Supplemental Figures for "A broadly-neutralizing antibody against *Ebolavirus* glycoprotein that potentiates the breadth and neutralization potency of other antibodies"

Figure S1.

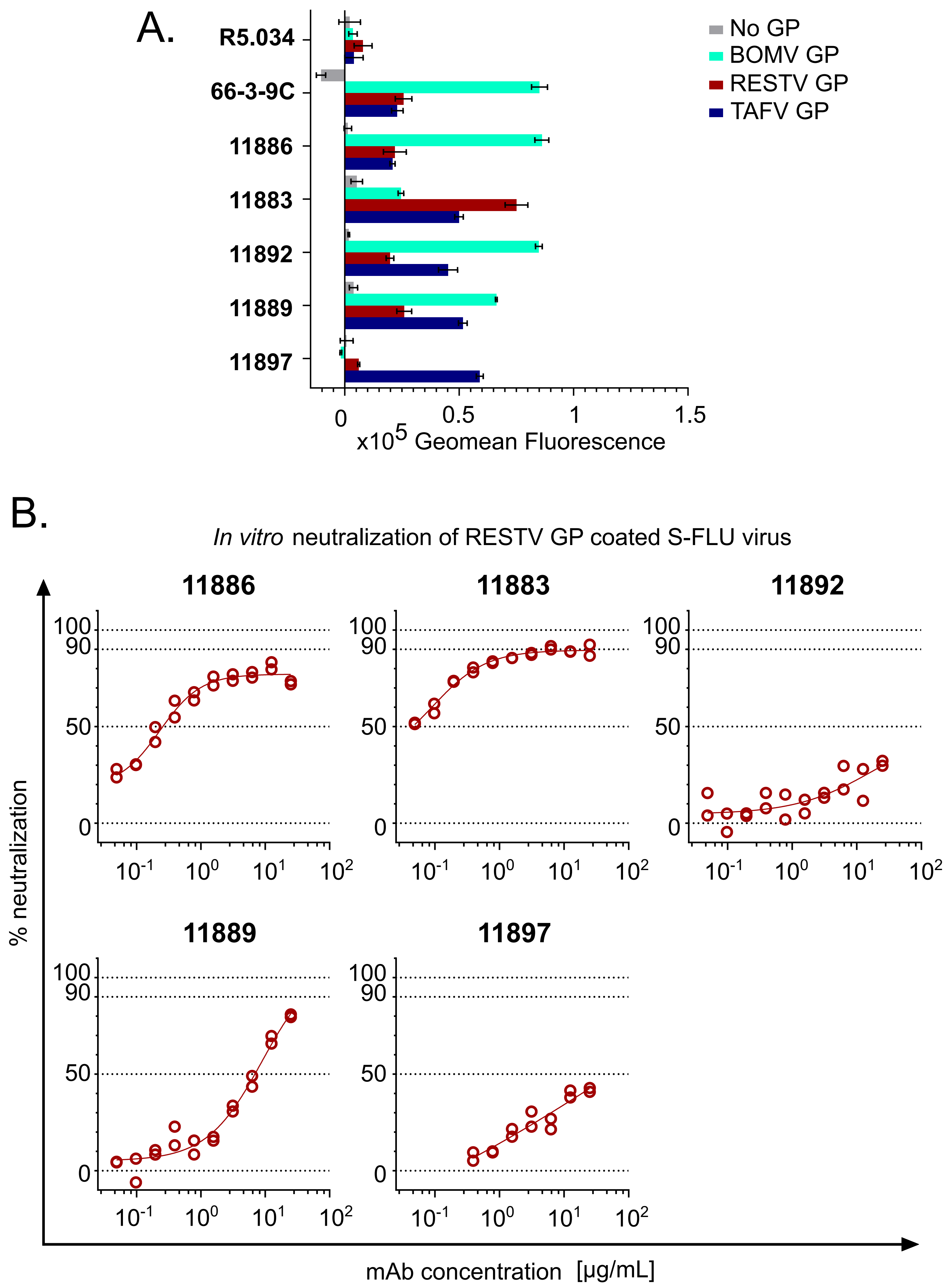
A.
R5.034
66-3-9C
11886
11883
11892
11889
11897
No GP
BOMV GP
RESTV GP
TAFV GP
0
0.5
1
1.5
x10^5 Geomean Fluorescence
B.
In vitro neutralization of RESTV GP coated S-FLU virus
11886
11883
11892
11889
11897
% neutralization
100
90
50
0
10^-1
10^0
10^1
10^2
mAb concentration [μg/mL]

Figure S2.

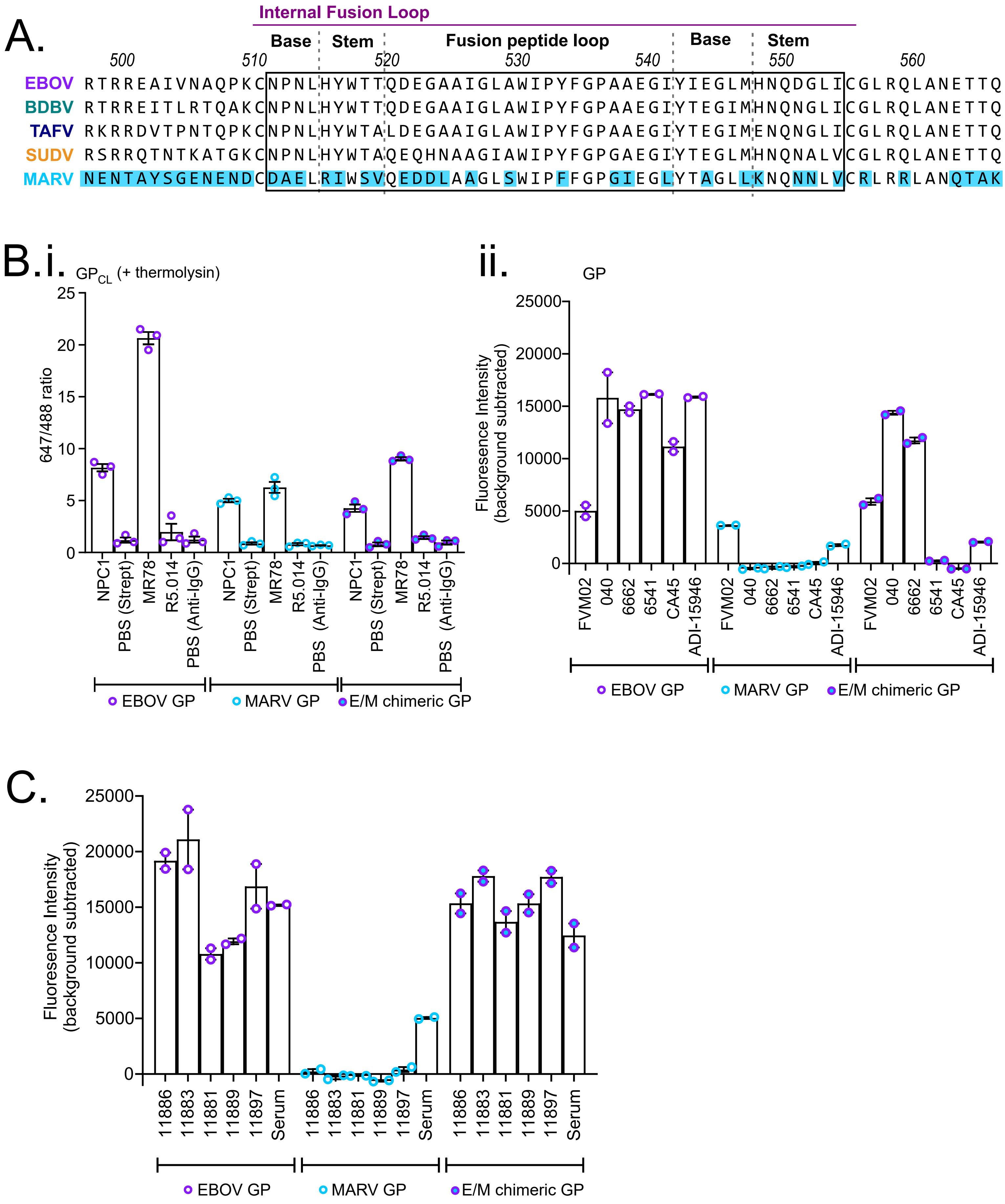
A.
Internal Fusion Loop
Base
Stem
Fusion peptide loop
Base
Stem
500
510
520
530
540
550
560
EBOV RTRREAIVNAQPKCNPNLHYWTTQDEGAAIGLAWIPYFGPAAEGIYIEGLMHNQDGLICGLRQLANETTQ
BDBV RTRREITLRTQAKCNPNLHYWTTQDEGAAIGLAWIPYFGPAAEGIYTEGIMHNQNGLICGLRQLANETTQ
TAFV RKRRDVTPNTQPKCNPNLHYWTALDEGAAIGLAWIPYFGPAAEGIYTEGIMENQNGLICGLRQLANETTQ
SUDV RSRRQTNTKATGKCNPNLHYWTAQEQHNAAGIAWIPYFGPGAEGIYTEGLMHNQNALVCGLRQLANETTQ
MARV NENTAYSGENENDCDAELRIWSVQEDDLAAGLSWIPFFGPGIEGLYTAGLLKNQNNLVCRLRRLANQTAK
B.i. $GP_{CL}$ (+ thermolysin)
647/488 ratio
25
20
15
10
5
0
NPC1
PBS (Strept)
MR78
R5.014
PBS (Anti-IgG)
EBOV GP
MARV GP
E/M chimeric GP
ii.
GP
Fluorescence Intensity (background subtracted)
25000
20000
15000
10000
5000
0
FVM02
040
6662
6541
CA45
ADI-15946
EBOV GP
MARV GP
E/M chimeric GP
C.
Fluoresence Intensity (background subtracted)
25000
20000
15000
10000
5000
0
11886
11883
11881
11889
11897
Serum
EBOV GP
MARV GP
E/M chimeric GP

Figure S3.

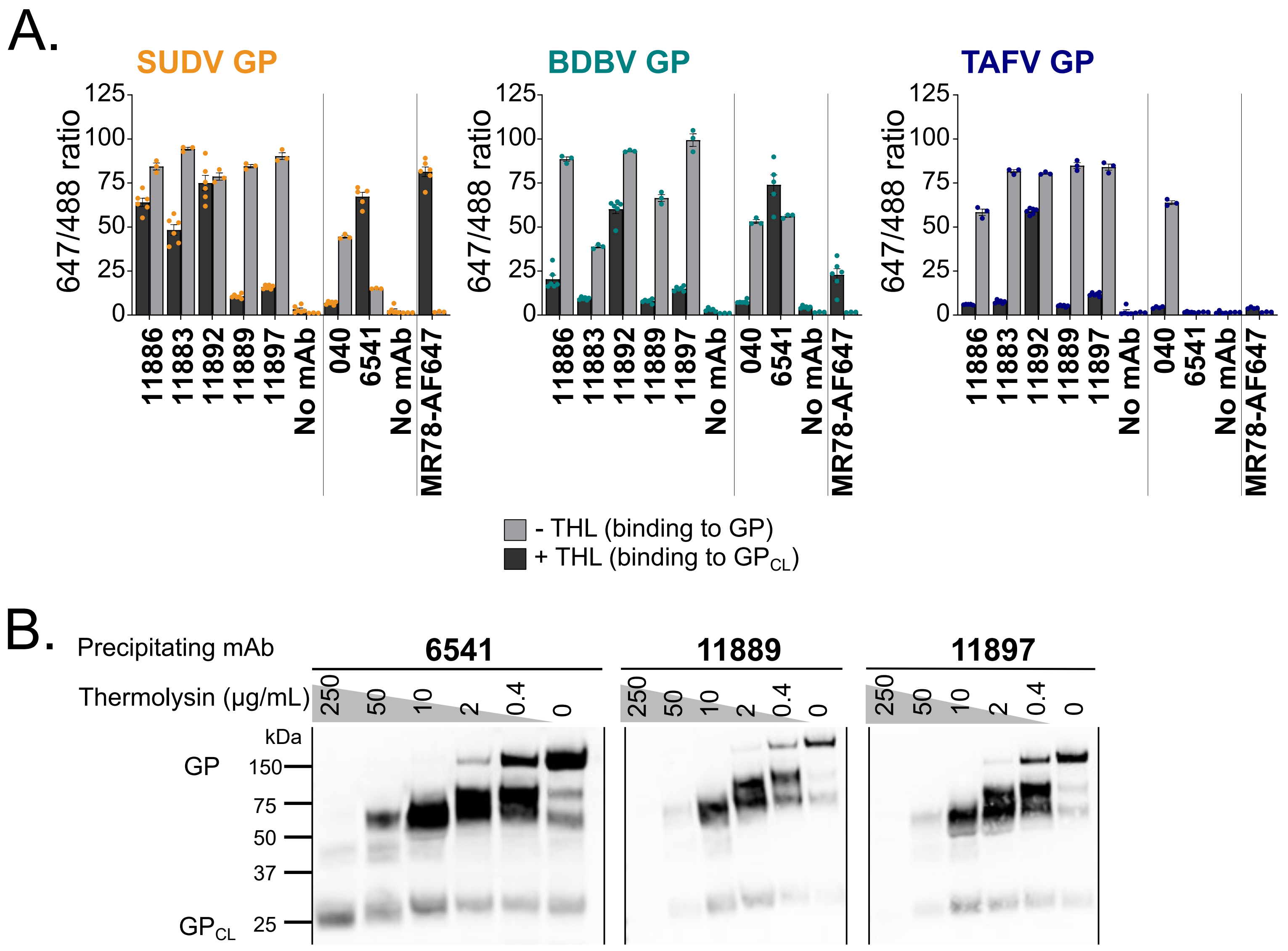
A.
SUDV GP
BDBV GP
TAFV GP
647/488 ratio
125
100
75
50
25
0
11886
11883
11892
11889
11897
No mAb
040
6541
No mAb
MR78-AF647
- THL (binding to GP)
+ THL (binding to GPCL)
B.
Precipitating mAb
6541
11889
11897
Thermolysin (µg/mL)
250
50
10
2
0.4
0
kDa
GP
150
75
50
37
GPCL
25

Figure S4.

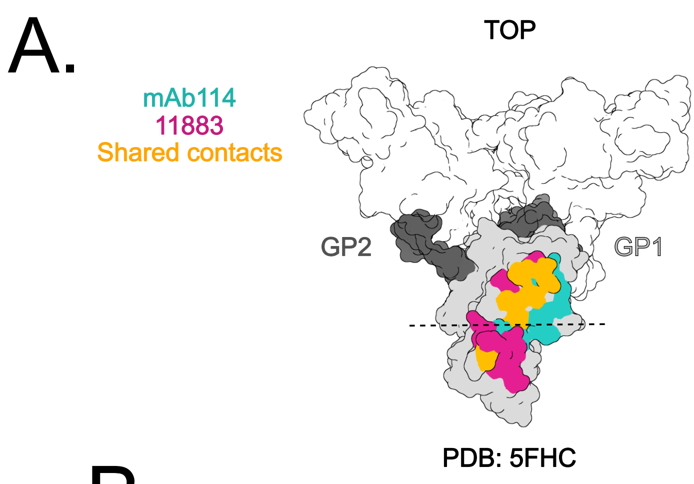


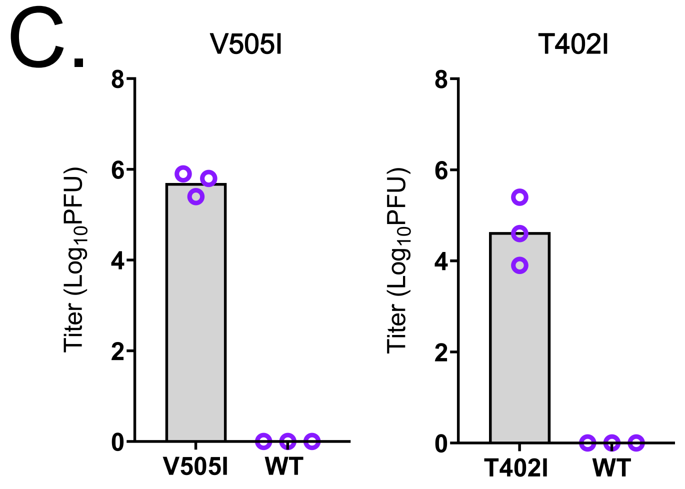


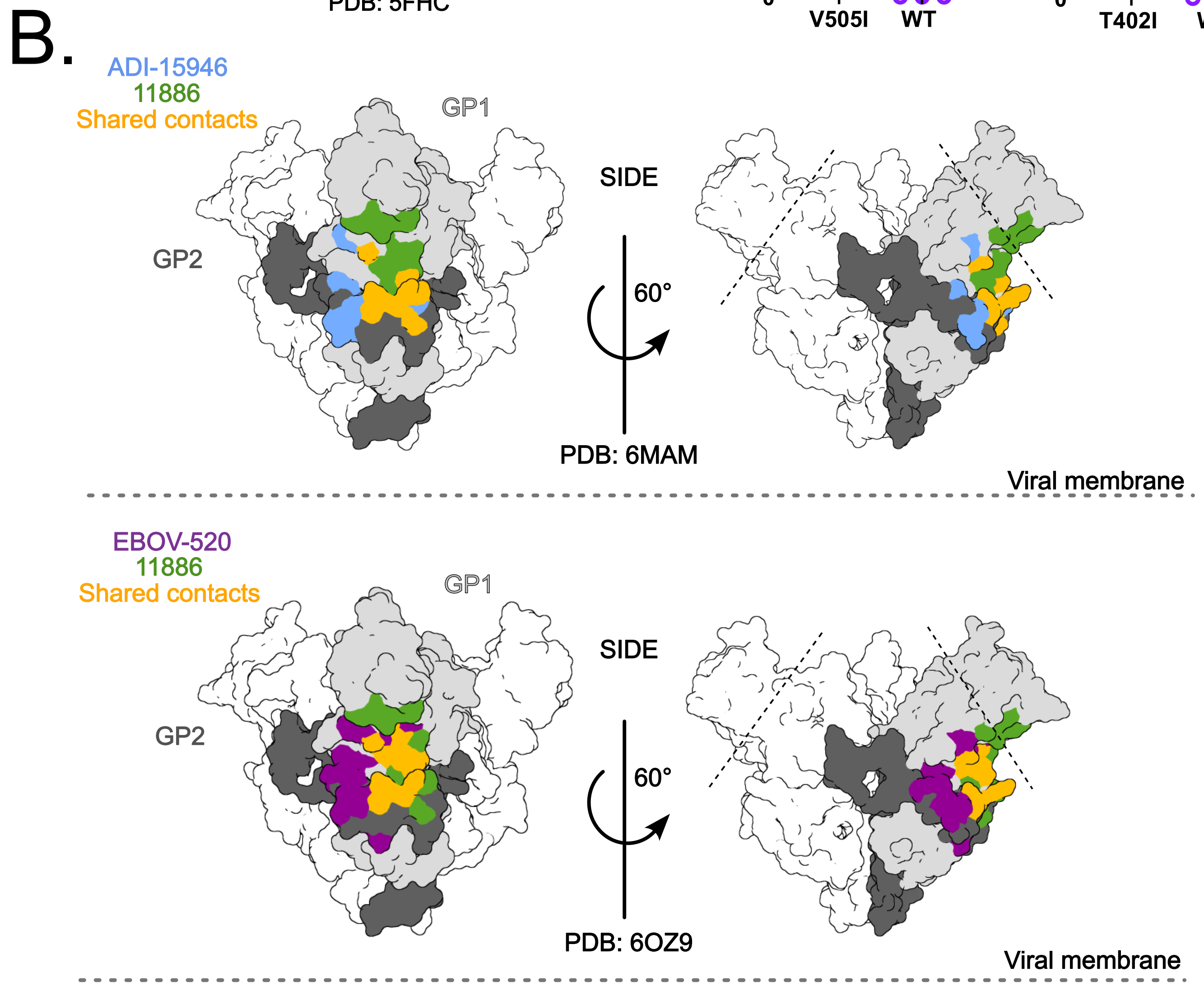


D.

```
                ...     |----:----|----:----|----:----|----:----|----:----|----:----|----:----|----:----|----:---          ...
Consensus    ...69     NLEGNGVATDVPSATKRWGFRXGVPPKVVNYEAGEWAENCYNLEIKKPDGSECLPEAPXGVRGFPRCRYVHKVSGTGPCPGXXAFHKEG   157...
EBOV         ...69     NLEGNGVATDVPSATKRWGFRSGVPPKVVNYEAGEWAENCYNLEIKKPDGSECLPAAPDGIRGFPRCRYVHKVSGTGPCAGDFAFHKEG   157...
BDBV         ...69     NLEGNGVATDVPTATKRWGFRAGVPPKVVNYEAGEWAENCYNLDIKKADGSECLPEAPEGVRGFPRCRYVHKVSGTGPCPEGYAFHKEG   157...
SUDV         ...69     NLEGSGVSTDIPSATKRWGFRSGVPPKVVSYEAGEWAENCYNLEIKKPDGSECLPPPPDGVRGFPRCRYVHKAQGTGPCPGDYAFHKDG   157...
TAFV         ...69     NLEGNGVATDVPTATKRWGFRAGVPPKVVNCEAGEWAENCYNLAIKKVDGSECLPEAPEGVRDFPRCRYVHKVSGTGPCPGGLAFHKEG   157...
RESTV        ...70     NLEGNGIATDVPSATKRWGFRSGVPPKVVSYEAGEWAENCYNLEIKKSDGSECLPLPPDGVRGFPRCRYVHKVQGTGPCPGDLAFHKNG   158...
BOMV         ...65     NLEGNGIATDVPSATKRWGFRAGVPPKVVGYEAGEWAENCYNLEIKKPDGSECLPMAPEGIRGFPRCRYVHKVSGTGSCESGFAFHKEG   153...

               ...     |----:----|----:----|----:----|----:----|----:----|----:----|----:---     ...     ----:----|--      ...
Consensus   ...219     XYEAXNFGTNEXXFLFKVDNLTYVQLEXRFTPQFLVQLNETIYTNGRRSNTTGRLIWKVNPTXDTDVGEW  288...501  XNTQXKCNPNLH   512...
EBOV        ...219     RYQATGFGTNETEYLFEVDNLTYVQLESRFTPQFLLQLNETIYTSGKRSNTTGKLIWKVNPEIDTTIGEW  288...505  VNAQPKCNPNLH   516...
BDBV        ...219     NYVADNFGTNMTNFLFQVDHLTYVQLEPRFTPQFLVQLNETIYTNGRRSNTTGTLIWKVNPTVDTGVGEW  288...505  LRTQAKCNPNLH   516...
SUDV        ...219     EYEIENFGAQHSTTLFKIDNNTFVRLDRPHTPQFLFQLNDTIHLHQQLSNTTGRLIWTLDANINADIGEW  288...505  TKATGKCNPNLH   516...
TAFV        ...219     NYVVDNFGTNTTEFLFQVDHLTYVQLEARFTPQFLVLLNETIYSDNRRSNTTGKLIWKINPTVDTSMGEW  288...505  PNTQPKCNPNLH   516...
RESTV       ...220     SYEMSNFGGNESNTLFKVDNHTYVQLDRPHTPQFLVQLNETLRRNNRLSNSTGRLTWTLDPKIEPDVGEW  289...506  QNTANKCNPDLY   517...
BOMV        ...215     EYETTRFGTNRSAFWFKVDNLTFVQLESRFTPQFLVELNETIYIEGKRSNTTGRLIWQVNSRVDTDVGEW  284...501  EIRGAKCNPNLH   512...
```

Figure S5.

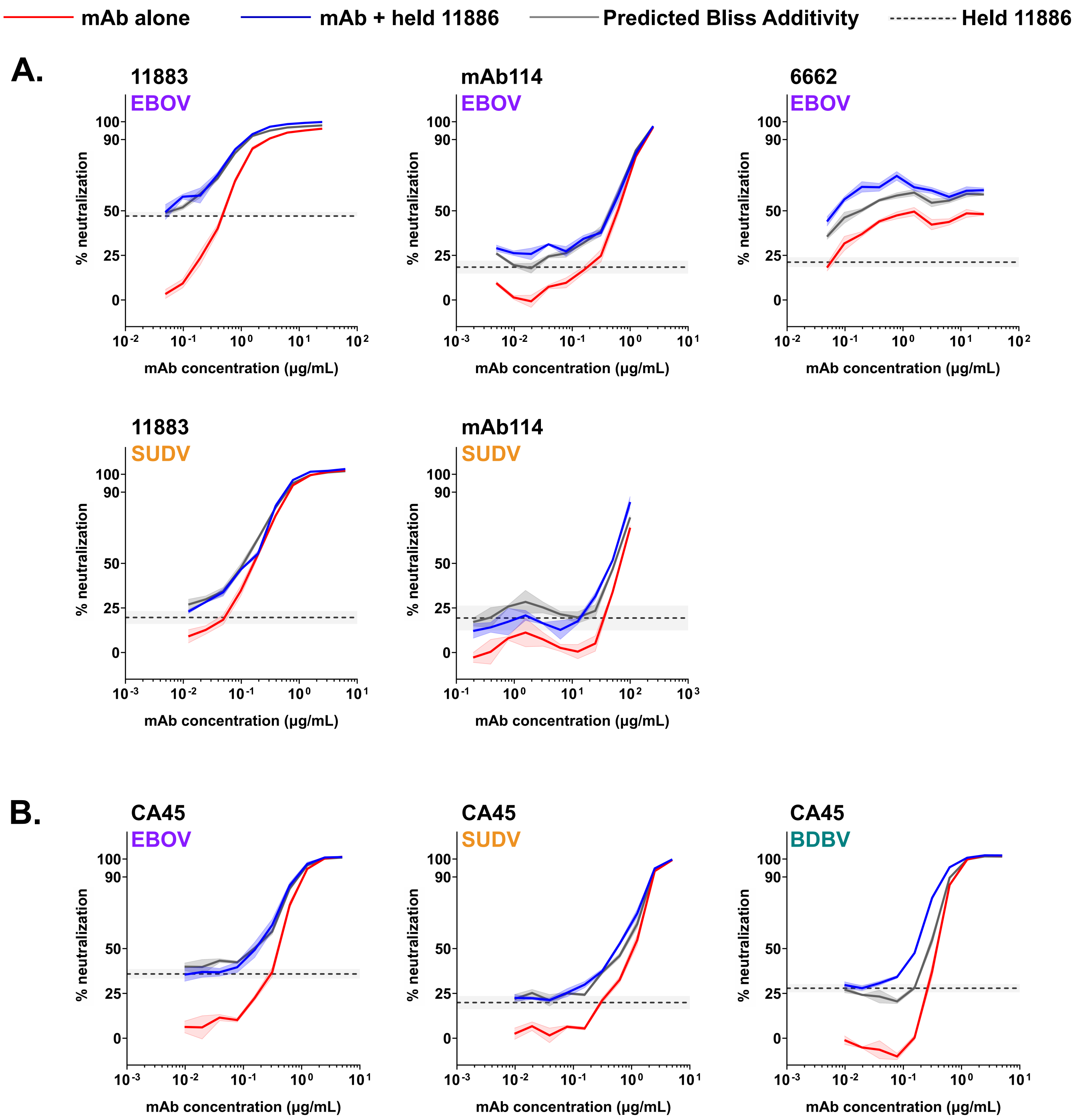
mAb alone
mAb + held 11886
Predicted Bliss Additivity
Held 11886
A.
11883
EBOV
mAb114
EBOV
6662
EBOV
11883
SUDV
mAb114
SUDV
B.
CA45
EBOV
CA45
SUDV
CA45
BDBV
% neutralization
mAb concentration (µg/mL)

Figure S6.

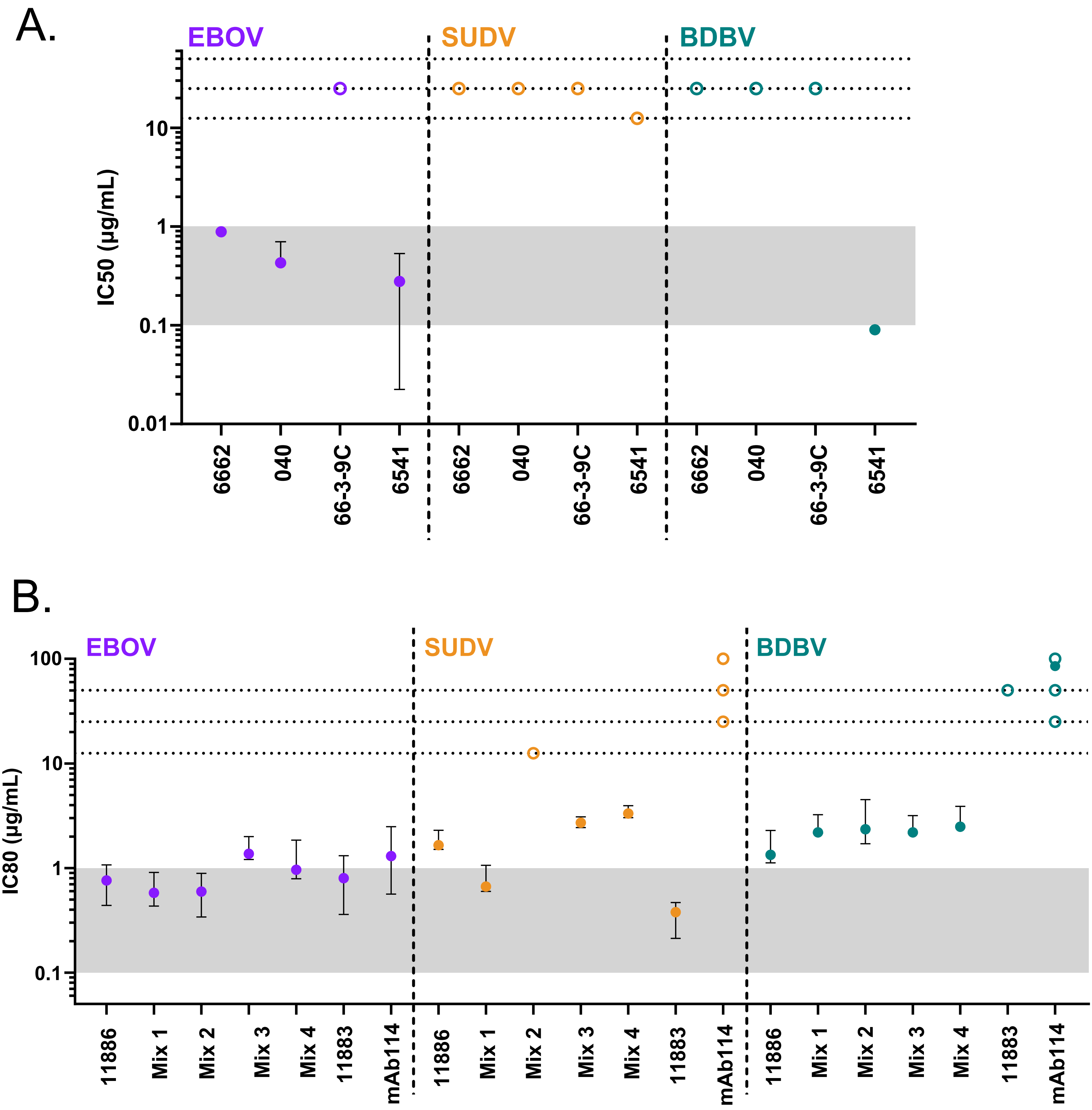


| | |
|---|---|
| Mix 1 | 11883 + 66-3-9C + 11886 |
| Mix 2 | 6662 + 040 + 66-3-9C + 6541 |
| Mix 3 | mAb114 + 66-3-9C + 11886 |
| Mix 4 | 6662 + 040 + 66-3-9C + 11886 |
