## Supplemental Figure Legends for "A broadly-neutralizing antibody against *Ebolavirus* glycoprotein that potentiates the breadth and neutralization potency of other antibodies"

### Supplemental Figures and Legends

*Supplemental Figure 1. Binding to TAFV, BOMV and RESTV GP and RESTV pseudovirus neutralization by rabbit mAb panel*

**A) Binding of broadly reactive rabbit mAb panel to full-length transmembrane GPs from *Ebolavirus* species that have not caused multiple outbreaks of EVD in humans.** MDCK SIAT-1 cells expressing TAFV GP (dark blue), RESTV GP (dark red), BOMV GP (turquoise) or parental cells not expressing GP (No GP, grey) were incubated with 50 µg/mL mAb, and mAb binding was detected using an Alexa Fluor 647 anti-IgG conjugate secondary antibody. R5.034 (non-anti-GP mAb) and 66-3-9C (broadly reactive anti-GP mAb) were included as negative and positive controls, respectively. Background (mean of 8 wells per assay plate incubated with relevant secondary only) was subtracted from data. Mean and SEM of duplicates shown for each mAb.

**B) *In vitro* neutralization of RESTV GP pseudotyped S-FLU virus.** S-FLU viruses coated with RESTV GP were incubated with mAb, and MDCK SIAT-1 cells then infected with the mAb-virus mixture. Percent neutralization of S-FLU virus by titrated mAb was calculated using maximal (no antibody added) and minimal fluorescence (no virus added) signals within each assay. Duplicates within assay for each mAb concentration and calculated non-linear regression curves are shown.

*Supplemental Figure 2. Development of chimeric GP for assessment of fusion loop binding.*

**(A) Alignment of Internal Fusion Loop (IFL) sequences from *Ebolaviruses* and Marburgvirus (MARV) GP sequences obtained from NCBI Reference database.** EBOV (Ebola virus/H.sapiens-wt/GIN/2014/Makona-C15; ATY51135), SUDV (Sudan virus/H.sapiens-tc/UGA/2000/Gulu-808892; YP_138523.1), BDBV (Bundibugyo virus/H.sapiens-tc/UGA/2007/Butalya-811250; YP_003815435.1), TAFV (Tai Forest virus/H.sapiens-tc/CIV/1994/Pauleoula-CI; YP_003815426.1), MARV (Marburg virus/H.sapiens-tc/KEN/1980/Mt.Elgon-Musoke; YP_001531156.1). Numbering associated with EBOV sequence. Region in box indicates sequences that are swapped to produce chimeric EBOV GP with

MARV IFL (“E/M chimeric GP”). Numbered according to EBOV sequence. Differences between MARV and EBOV sequences are highlighted in blue in the MARV sequence.

**(B) Validation of chimeric GP as a tool to assess IFL binding**. (i) Confluent monolayers of MDCK-SIAT 1 cells expressing the GPs on their surface were treated with thermolysin to reveal the RBR, then incubated with 10 µg/mL biotinylated NPC1, MR78 (pan-filovirus RBR-binding mAb control) or R5.014 (irrelevant malaria mAb control) in triplicate. NPC1 was detected using a streptavidin Alexa Fluor 647 conjugate; MR78 and R5.014 were detected using anti-human IgG Alexa Fluor 647 conjugate. Cells were then stained with wheat germ agglutinin Alexa Fluor 488 conjugate to normalize for number of cells remaining in each well after thermolysin treatment. The mean and SEM of the ratio between fluorescence intensities of the 647 and 488 conjugates of triplicates in each assay are shown. (ii) Confluent cell monolayers were incubated with reference mAbs at 10 µg/mL which were then detected using anti-human IgG AlexaFluor 488 conjugate. The mean and SEM of duplicates in each assay are shown. Representative data are shown for mAbs 040, 6662, 6541 and FVM02 which were repeated in N=3 experiments.

**(C). Broadly reactive rabbit mAbs do not lose binding to chimeric GP with MARV IFL.** Confluent monolayers of MDCK-SIAT 1 cells expressing the GPs on their surface were incubated with test mAbs at 10 µg/mL, and mAb binding was detected using anti-rabbit IgG Alexa Fluor 488 conjugate. The mean and SEM of duplicates within each assay are shown. Representative data shown from N=2 repeats of each experiment. Serum from the rabbit immunized with *Ebolavirus* GPs for initial antibody discovery was included as a positive control.

*Supplemental Figure 3. Binding of mAb panel to cleaved and uncleaved Ebolavirus GPs*

**(A) Antibody binding to thermolysin (THL) cleaved cell surface-expressed SUDV, BDBV and TAFV GP was assessed in an immunofluorescence assay.** For each test mAb, binding to GP was tested under two conditions: GP-expressing cells were pre-incubated with THL to produce $GP_{CL}$, then incubated with test mAbs (dark grey); and mAbs were incubated with GP-expressing cells without THL (mid-grey).

mAb binding was then detected using Alexa Fluor 647 conjugate and cells stained with wheat germ agglutinin Alexa Fluor 488 conjugate. Mean and SEM for replicate wells within the assay (6 wells for GP with THL, 3 wells for GP without THL treatment) are shown. 040 (GC binding mAb) and 6541 (base binding mAb) are comparator mAbs showing THL sensitive and insensitive binding profiles. MR78 can only bind to EBOV GP when the GL and MLD are removed exposing the RBR [1], and is included as a control to confirm THL digestion of GP in the assay.

**(B) Immunoprecipitation of thermolysin cleaved EBOV GP by mAbs 6541, 11889 and 11897.** GP-expressing cells were treated to biotinylate surface proteins, then incubated with increasing concentrations of THL. Cells were washed then lysed. GP was immunoprecipitated from cell lysates using Protein A Sepharose and anti-GP mAb of interest. Samples were run on reducing SDS-PAGE, and bands were revealed using streptavidin Alexa Fluor 647 conjugate. Band at ~150 kDa is full-length GP; band at ~25 kDa is $GP_{CL}$ (GP1 core and GP2); other bands are products of sequential cleavage of MLD and GC. 6541 is a known base-binding mAb that can immunoprecipitate the GP1 core/GP2.

*Supplemental Figure 4. Comparison of 11883 and 11886 footprints with previously described antibodies.*

**(A) Comparison of 11883 footprint with mAb114.** GPΔmuc trimer ectodomain with 11883 contacts on the GP (pink) overlaid with mAb114 (teal) (PDB: 5FHC) contacts. Residues in common between the two footprints are shown in yellow.

**(B) Comparison of 11886 footprint with other $3_{10}$ pocket binding mAbs.** GPΔmuc trimer ectodomain with 11886 contacts on the GP (green) overlaid with ADI-14956 contacts (light blue) (PDB: 6MAM) or EBOV-520 (purple) (PDB: 6OZ9) contacts. Residues in common between the two footprints are shown in yellow. For monomer with footprints shown, GP1 is shaded in light grey and GP2 in dark grey. Dotted line indicates domains cleaved after cathepsin or THL treatment to generate $GP_{CL}$. In PDB: 6OZ9, the GP does not contain the GC, therefore any contacts between EBOV-520 and the GC are not modelled. However, in an additional structural study, PDB: 6PCI, a complex of EBOV GPΔmuc trimer with EBOV-

520 and a partner mAb that remodels the glycan cap, suggests that EBOV-520 does contact the GP1 in that structure, with contacts reported as Pro250, Gln251 and Gln255 with the α2 helix displaced upwards compared to the unliganded GP [2,3].

**(C) VPΔ30 EBOV with V505I or T402I mutations in the GP permits viral escape from neutralization by mAb 11886.** PFU; Plaque-forming units. WT; wild-type. Viruses incubated with 10 µg/mL 11886 mAb.

**(D) Alignment of *Ebolavirus* GP sequences showing conservation of 11886 and 11883 contact residues.** mAb contact residues on EBOV GP (and conserved residues in other GPs) are highlighted. 11886 (green) and 11883 (pink). Location of residue Val505 highlighted by black arrow.

*Supplemental Figure 5. Additive interactions between 11886 and other broadly reactive partner mAbs across Ebolavirus species in a pseudovirus neutralization assay.*

**(A) 11886 with partner RBR mAbs 11883, mAb114 and 6662.**

**(B) 11886 with partner fusion loop mAb CA45.**

For all data, a representative experiment of at least two independent repeats has been shown. Dotted line represents the mean inhibition given by a held concentration of 11886 with 95 % confidence limits shown in shaded grey. Solid lines represent mean of triplicate wells within an experiment, with shaded areas indicating the standard error. Red lines indicate the inhibition given by a titration series of the partner mAb alone. Blue lines indicate the inhibition given by a titration series of the partner mAb plus the held concentration of mAb 11886. Grey line represents the calculated Bliss Additivity value which assumes a purely independent and additive interaction between the partner mAb and 11886. Held concentration of 11886 ranged from 0.2-0.6 µg/mL between experiments to achieve target 20-45 % inhibition in a given assay.

*Supplemental Figure 6. Neutralization by component antibodies of cocktail mixes.*

Antibodies were tested for neutralization of EBOV (purple), SUDV (orange) and BDBV (teal) GP coated S-FLU pseudoviruses. Open symbols represent results where 50 % neutralization of virus was not achieved at the highest concentration of the test antibody assayed in that experiment. Horizontal dotted lines denote 12.5, 25 and 50 µg/mL of total antibody respectively.

**(A)** Median and 95 % CI of IC50 values of individual mAbs that were included in cocktail mixes (**Figure 7**) from N=2 to N=5 experiments are shown.

**(B)** Median and 95% CI of the IC80 values (concentration of antibody required to achieve 80 % virus neutralization) for antibody mixes and 11886 alone were calculated from N=3 experiments. For comparison, median and 95 % CI of IC80 values of individual antibodies 11883 and mAb114 calculated from N=2 to N=5 experiments run separately are also shown.
